# Plant A20/AN1 proteins coordinate different immune responses including RNAi pathway for antiviral immunity

**DOI:** 10.1101/622696

**Authors:** Li Chang, Ho-Hsiung Chang, Yi-Shu Chiu, Jui-Che Chang, Duen-Wei Hsu, Yuh Tzean, An-Po Cheng, Hsiang-Chia Lu, Hsin-Hung Yeh

**Affiliations:** Molecular and Biological Agricultural Sciences Program, Taiwan International Graduate Program, National Chung-Hsing University, Academia Sinica, Taiwan; Agricultural Biotechnology Research Center, Academia Sinica, Taipei, Taiwan; Graduate Institute of Biotechnology, National Chung-Hsng University, Taichung, Taiwan; Department of Biotechnology, National Kaohsiung Normal University, Kaohsiung, Taiwan; Department of Plant Pathology and Microbiology, National Taiwan University, Taipei, Taiwan

## Abstract

Salicylic acid (SA)-mediated immunity plays important roles in combating virus in plants. Two plant stress associated protein (SAPs) containing dual A20/AN1 zinc-finger domain were found to play important roles in SA-mediated immunity; however, detailed mechanisms remain elusive. In this study, another orchid homolog gene of *Pha13*, *Pha21*, was analyzed. Pha21 confers antiviral immunity in both transgenic orchid and *Arabidopsis* overexpressing Pha21. Expression of *Pha21* is early-induced by SA treatment, and is involved in the expression of the orchid homolog of the master regulator *NPR1*. Pha21 but not Pha13 is involved in the expression of key RNAi-related genes, *Dicer-like nuclease 4* (*DCL4*) and *Argonaut 1* (*AGO1*) in orchids. The involvement of SAPs in expression of orchid *DCL4* and *AGO1* is not limited to orchid, as AtSAP5 also plays essential role in the expression of *Arabidopsis DCL4* and *AGO1*. However, unlike Pha13 and AtSAP5, Pha21 does not play positive role in the expression of orchid homolog gene of *RNA-dependent RNA polymerase 1* (*RdR1*), an important gene in RNAi pathway. Pha21 can be found localized in the nucleus, and confers self-E3 ligase and ubiquitin binding activities. Functional domain analysis revealed that both A20 and AN1 domains of Pha21 are required for decreasing virus accumulation, and the AN1 domain plays a more important role in the expression of orchid *DCL4*. Collectively, our data suggests SA regulated SAPs play important roles in antiviral immunity and is involved in delicate regulation of key genes in RNAi-mediated pathway.

**IMPORTANCE:** Salicylic acid (SA)-mediated antiviral immunity plays an important role to protect plants from virus infection; however, the detailed mechanisms remain elusive. We previously demonstrated that two plant A20/AN1 proteins, orchid Pha13 and *Arabidopsis* AtSAP5, function similarly and serve as an important hub to regulate SA-mediated antiviral immunity. In this study, we identified another orchid A20/AN1 protein, Pha21, which is involved in SA-mediated antiviral immunity. In contrast to Pha13 and AtSAP5, Pha21 plays minor negative roles in the expression of *PhaRdR1* (orchid homolog of *RNA-dependent RNA polymerase 1*). However, Pha21 and AtSAP5, but not Pha13, are involved in the expression of important players in RNAi pathway, *Dicer-like nuclease 4* (*DCL4*) and *Argonaut 1* (*AGO1*), in orchid and *Arabidopsis*. Our data demonstrates that plant A20/AN1 proteins are conserved players in SA-mediated antiviral resistance among plants, and provide links between the A20/AN1 proteins and the RNAi pathway.

## INTRODUCTION

The plant hormone salicylic acid (SA) is important to trigger plant immunity especially against biotrophic pathogens such as viruses (1–4). In facing pathogen invasion, plants recognize conserved microbe-associated molecular patterns (MAMPs) by pattern-recognition receptors and trigger pathogen (or pattern)-triggered immunity (PTI) as the first line of defense (5, 6). Although PTI provides protection from invasion by most pathogens, some have evolved, and can utilize different effectors to suppress PTI for successful infection (7). Plants have evolved resistance (R) proteins capable of detecting these effectors to trigger effecter-triggered immunity (ETI) for a second line of plant defense (7). ETI is generally associated with programmed cell death leading to the formation of necrotic lesions, which is also known as hypersensitive response (HR) to prevent the further spreading of pathogen infection (1).

Current evidence suggests that the PTI is important to limit virus infection (8–10), and the viral double strand RNA (dsRNA) has been shown to serve as a conserved MAMP (10). In addition, ETI also plays an important role in combating virus infection in plants. For example, the N gene from *Nicotiana glutinosa* serves as an R protein to specifically recognize the effector from *Tobacco mosaic virus* (TMV) and trigger the ETI (11–14). SA plays important roles in triggering PTI and ETI (15). In addition, the TMV-infected tobacco plant showing local necrotic lesions was found to become more resistant against secondary virus infection in the distal leaves (16, 17). This systemic immune response is known as systemic acquired resistance, which is a common plant immune response and plays an important role in protecting plants from pathogen infection (4).

In addition, RNA silencing or RNA interference (RNAi) has also been demonstrated to play an important role in combating virus infection (18, 19). RNAi is activated through the appearance of the viral dsRNA upon virus infection. The dsRNA can be cleaved to short small-interfering dsRNA (siRNA) of 21–24 nucleotides (nt) in size by the Dicer-like (DCL) nuclease (20). DCL2 and DCL4 in *Arabidopsis* play an important role in generating the virus-derived siRNA (vsiRNA) (19, 21, 22). The vsiRNA is further unwound and one strand (guide-strand) is incorporated into the RNA-induced silencing complex (RISC) (19). This complex targets viral RNA with a sequence that is complementary to the guide-strand and degrades the target RNA by the catalytic component of the RISC, Argonaut (AGO) nucleases (19, 20, 23, 24). Among the *Arabidopsis* AGOs, AGO1 plays an important role in the antiviral immunity against RNA viruses. (24). More vsiRNA can be further generated *de novo* through the cellular RNA-dependent RNA polymerases (RDRs) to trigger the secondary RNA silencing against viruses (25–27). SA also plays important roles in the RNAi-pathway as SA treatment can induce genes involved in the RNAi pathway including *RDRs* in *Arabidopsis*, *Nicotiana tabacum and* Tomato, and *DCLs* and *AGOs* in Tomato (28–32).

SA induces the fluctuation of redox and serves as a signal to activate sets of defense genes (33, 34). The SA-induced redox change can modify the NPR1 (nonexpressor of pathogenesis-related protein 1) from the multimeric protein complex to a monomer via the oxidoreductases, Thioredoxin-3 and Thioredoxin-5 (35, 36). The NPR1 monomer moves into the nucleus and activates multiple defense-related genes including the *PR* (*pathogenesis-related*) gene in the SA-signaling pathway (37–39). In addition to the NPR1-dependent immune pathway, some data also suggest that the existence of a NPR1-independent pathway contributes to the virus resistance (40, 41). Despite the importance of the SA governed immune pathway in antiviral immunity, the regulation of this pathway remains largely elusive.

Proteins containing zinc-finger A20 and/or AN1 domains play an important role in plant response against various abiotic stresses, and are known as stress associated protein (SAPs). SAPs are conserved among different organisms (42, 43). Different numbers of SAP homologs (range from 1 to 19) have been identified in organisms including protists, fungi, animals, plants, and humans (42, 43). Compared to animals, more of these proteins are found in plants. So far, 18, 14 and 14 SAPs have been identified in rice, *Arabidopsis*, and *Phalaenopsis* orchid, respectively (42, 44). Biochemical studies of proteins containing A20 and/or AN1 domains revealed that the A20 domain confers E3 ligase and ubiquitin binding activity (43–49). The A20 domain of human A20, Rabex-5 (guanine nucleotide exchange factor), *Arabidopsis* AtSAP5 and orchid Pha13 have been reported to exhibit E3 ligase activity (44, 45, 49, 50). In addition, it has been shown that the A20 domain can also bind to various ubiquitin chains (44, 46–48). In contrast to the A20 domain, the biochemical function of the AN1 domain is not fully understood.

Our recent study indicates that plant SAPs play a pivotal role in SA governed antiviral immunity (44). Orchid *Pha13* and *Arabidopsis AtSAP5* are induced by SA treatment at the early stage, and are involved in expression of orchid or *Arabidopsis NPR1* and *NPR1*-independent immune responsive genes including the induction of plant *RdR1*. In addition to Pha13, our previous virus-induced gene silencing (VIGS) assay also allowed us to identify the involvement of orchid *PhaTF21* (designated here as *Pha21*) in SA–regulated immune response genes (51). Pha13 and Pha21 share 69.5% amino sequence identity, and are most closely related among the orchid SAPs (44). Whether Pha21 and Pha13 work in a cooperative manner in plant antiviral immunity remains to be explored.

Here, we performed a detailed study of Pha21, and compared its biochemical and physiological function to Pha13. Pha21 and Pha13 share similar biochemical properties including E3-ligase and ubiquitin binding activities. Our results also indicate that similar to *Pha13*, *Pha21* is early-induced by SA, involved in the expression of the *PhaNPR1* and *PhaNPR1*-independent genes, and plays an important role in antiviral immunity. Importantly, our studies show that Pha21 is involved in expression of orchid *DCL4* and *AGO1*. Together with our previous data, these studies suggest that Pha13 and Pha21 coordinate the expression of genes important in the RNAi pathway including orchid *RdR1*, *DCL4* and *AGO1*. In addition, our previous data and data presented in this study also showed that *Arabidopsis* AtSAP5 is involved in expression of *RdR1*, *DCL4* and *AGO1*. Collectively, our data indicates that plant A20/AN1 proteins involved antiviral immunity are conserved among plants, and the antiviral immunity is partly through the RNAi pathway. Our findings suggest that A20/AN1 proteins may serve as a link between SA and the RNAi pathway at the early stage of SA induced-antiviral immunity.

## RESULTS

### Sequence and expression pattern analysis of Pha21

Previously, we reported that Pha21 (Orchidstra 2.0 database, http://orchidstra2.abrc.sinica.edu.tw, accession number PATC144963) is involved in the SA induced immune pathway in *P. aphrodite* (52). Pha21 contains dual zinc-finger domains, A20 and AN1, in the N-terminal and C-terminal, respectively (Fig. 1A-C), and belongs to the stress associated protein (SAP) family (43). Among the SAPs of *P. aphrodite*, Pha21 is most related to our previously reported Pha13 and shared 69.5% amino acid sequence identity (44). The A20 and AN1 zinc finger domains of Pha21 also shared high amino acid sequence identity among SAPs from different species including AtSAP5 from *Arabidopsis,* and OsSAP3 and OsSAP5 from *Oryza sativa* (Fig. 1B). (44). Unlike Pha13, no nuclear localization signal was identified in Pha21 (Fig. 1A).

**FIG 1.**
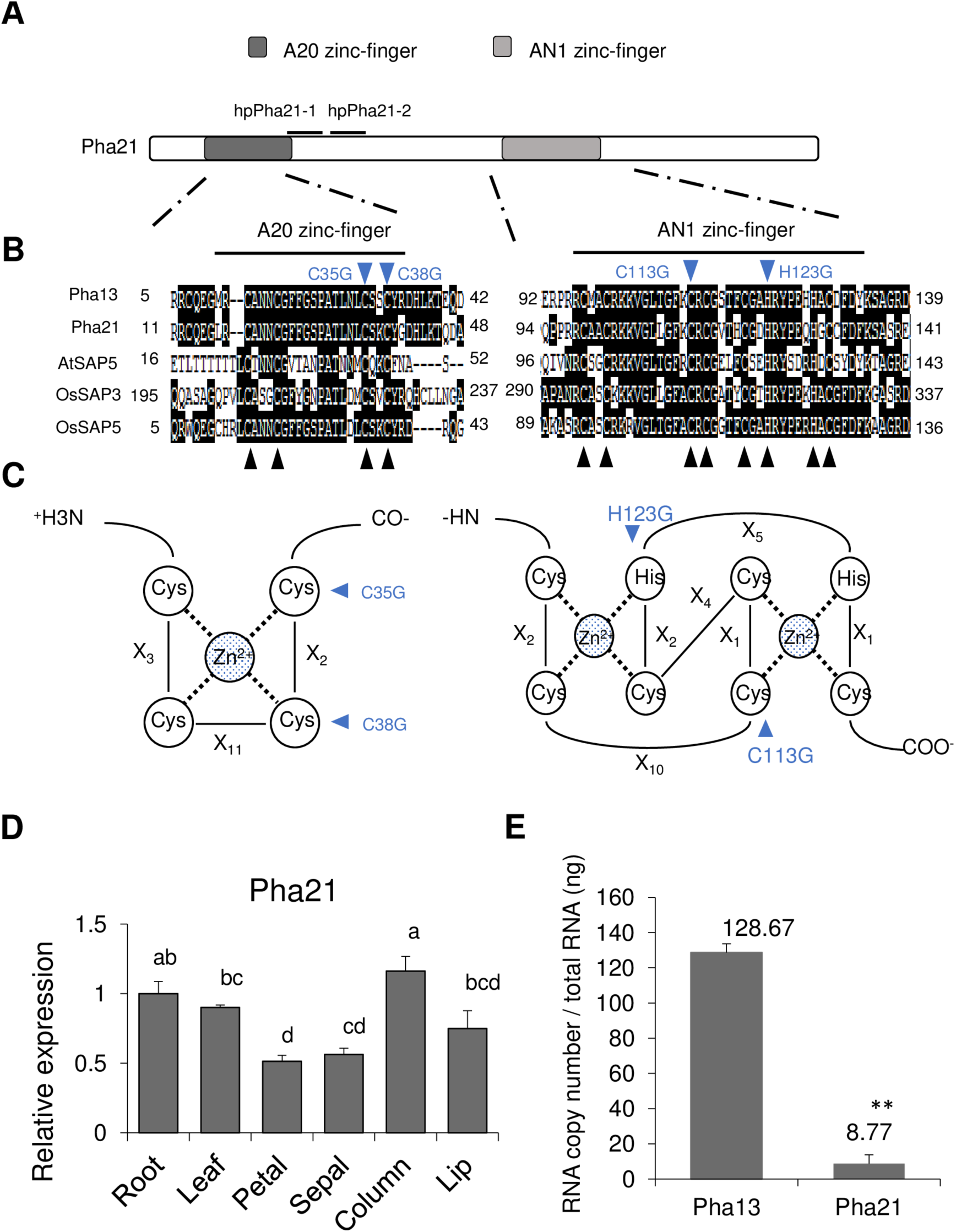
Sequence and expression analysis of Pha21 (A) A schematic representation of domain organization of Pha21. The open rectangle indicates the entire protein. A20 (dark grey rectangle) and AN1 (light grey rectangle) and 21-nucleotide position (short black line) used for designing hairpin RNA of Pha21 (hpPha21-1 and hpPha21-2) are indicated. (B) Amino sequence alignment of A20/AN1 zinc finger domains of Pha21 with stress-associated proteins from *Phalaenopsis aphrodite* (Pha13, accession number: PATC148746), *Arabidopsis thaliana* (AtSAP5, accession number: AT3G12630), and *Oryza sativa* (OsSAP3, accession number: LOC_Os01g56040.1; OsSAP5, accession number: LOC_Os02g32840.1) is shown. The black box indicates the conserved amino acid sequences. The black triangle indicates the conserved cysteine (C) and histidine (H) residues. (C) Primary sequence organization of A20 and AN1 domains. Xn: the number of amino acid residues between zinc ligands. (B-C) The mutation positions for domain functional analysis are indicated with a blue triangle. (D) Expression level of *Pha21* was analyzed by qRT-PCR from the root, leaf, petal, sepal, column, and lip of *P. aphrodite*. The RNA level of the root was set to 1. Data represent mean ± SD; n = 3 technical replicates; different letters indicate statistically significant differences analyzed by one-way analysis of variance (ANOVA) Tukey’s test (*P* < 0.05). The experiment was repeated at least three times with similar results, and one representative experiment is shown. *PhaUbiquitin 10* was used as an internal control for normalization. (E) Absolute quantitative analysis of *Pha13* and *Pha21* expression. The mRNA copy numbers of Pha13 and Pha21 in leaves of *P. aphrodite* were absolutely quantified by droplet digital PCR. Total RNA (ng) was used for normalization. Data represent mean ± SD; n = 3 biological replicates; **, *P* < 0.01, Student’s t test compared Pha13 with Pha21

The expression of *Pha21* was analyzed in different tissues of *P. aphrodite* orchid including root, leaf, septal, petal, lip, and column. The results revealed that *Pha21* expression was higher in the column, root and leaf (Fig. 1D). In addition, we also analyzed the absolute expression of *Pha21* and *Pha13* in the leaves of *P. aphrodite* by use of droplet digital PCR. The results indicated that *Pha21* RNA expression is about 15 times lower than *Pha13* in leaves of *P. aphrodite* (Fig. 1E)

### Pha21 is involved in the expression of SA responsive genes, *PhaNPR1* and *PhaPR1*

To further analyze the role of *Pha21* in the SA-induced immune pathway, we transiently expressed two hairpin RNA (will generate 21 nt siRNA) of Pha21 (35S::hpPha21-1 and 35S::hpPha21-2) separately to knockdown Pha21 by agroinfiltration in *P. aphrodite* carrying hairpin RNA expression construct, phpPha21-1 and phpPha21-2 (Fig. 2A). Samples collected from the infiltrated site of orchid leaves were used to detect the RNA level of Pha21, PhaNPR1, and PhaPR1 by quantitative RT-PCR (qRT-PCR). As shown in Fig. 2A, the expression of *PhaNPR1* and *PhaPR1* was decreased in both Pha21-silenced plants. In addition to transient knockdown of Pha21, we also transiently overexpressed Pha21 (35S::FLAG-Pha21) through agroinfiltration in the leaves of *P. aphrodite.* However, transient overexpression of Pha21 in leaves did not significantly affect the RNA level of PhaNPR1 and PhaPR1 (Fig. 2B).

**FIG 2.**
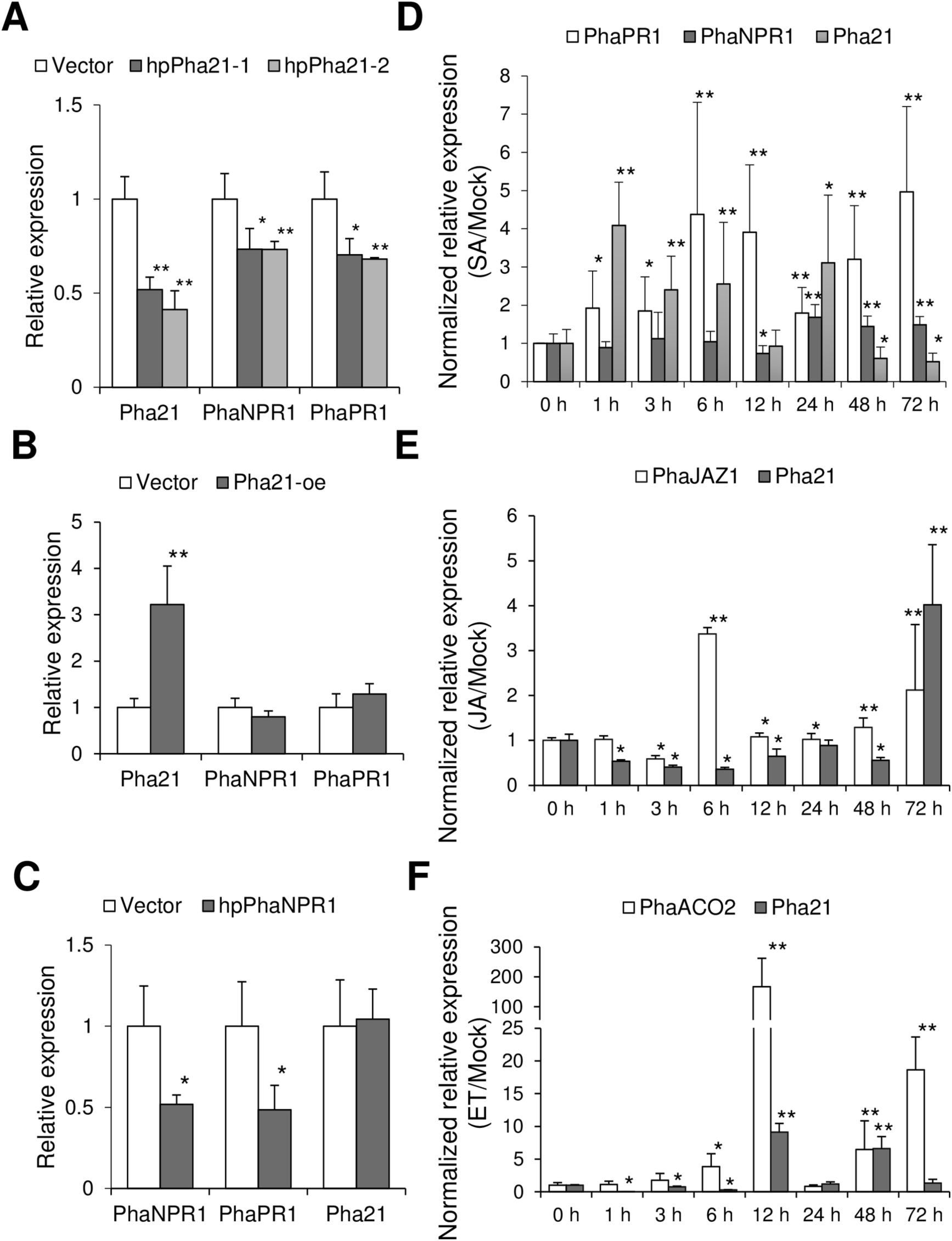
Pha21 is involved in the expression of *PhaPR1* and *PhaNPR1,* and responds to phytohormone treatment. (A-B) qRT-PCR was performed to quantify the expression level of *Pha21*, *PhaNPR1*, and *PhaPR1* from leaves of *P. aphrodite* infiltrated with agrobacterium carrying control vector (Vector); hairpin RNA vector to knock down Pha21 (hpPha21-1 and hpPha21-2; A); and overexpression vector of Pha21 (Pha21-oe; B). The expression level of plants infiltrated with control vector was set to 1. (C) Transient silencing of PhaNPR1. qRT-PCR was performed to quantify the expression level of *PhaNPR1*, *PhaPR1*, and *Pha21* from leaves of *P. aphrodite* infiltrated with agrobacterium carrying control vector (Vector) or hairpin RNA (hpRNA) vector to knock down PhaNPR1 (hpPhaNPR1). The expression level of plants infiltrated with control vector was set to 1. (A-C) Data represent mean ± SD; n = 3 biological replicates; **, *P* < 0.01; *, *P* < 0.05, Student’s t test compared to Vector. (D-F) Time-course expression of *Pha21* under different plant hormone treatments in *P. aphrodite*. qRT-PCR was performed to analyze the expression level of *Pha21* from leaves treated with SA (D), JA (E), and ET (F) at different hours (h) post-treatment. Buffer treatment was used as a mock control. Results of qRT-PCR were relative to that of mock at individual time courses for relative quantification. The expression level at 0 h was set to 1 for comparison between different time courses. *PhaPR1* and *PhaNPR1* were SA-responsive marker genes. *PhaJAZ1* and *PhaACO2* were JA and ET–responsive marker genes, respectively. Data represent mean ± SD; n = 3 biological replicates; **, *P* < 0.01; *, *P* < 0.05, Student’s t test compared to 0 h. (A-F) *PhaUbiquitin 10* was used as an internal control for normalization.

In addition, we also transiently silenced PhaNPR1 (phpPhaNPR1) in *P. aphrodite* to analyze the expression of *Pha21*. The results showed that the silenced PhaNPR1 showed decreased expression of *PhaPR1*; however, no obvious effect was observed on the expression of *Pha21* (Fig. 2C). These results suggest that Pha21 is involved in the expression of *PhaNPR1*, but not vice versa.

### *Pha21* is induced by SA, jasmonic acid and ethylene

We also tested whether *Pha21* was induced by defense-related plant hormones including SA, jasmonic acid (JA) and ethylene (ET). Orchid plants were treated with SA, JA and ET and samples were collected after treatment at different time points (up to 72 h) for analysis of the expression of *Pha21* and the marker genes of each plant hormone by qRT-PCR. The induction of *Pha21* by SA was observed at 1 h post-treatment (Fig. 2D). In addition, *Pha21* was also induced by JA and ET at 72 h and 12 h, respectively (Fig. 2E and F). Our results showed that *Pha21* can respond to multiple defense-related plant hormones, and SA induced *Pha21* expression at the very early stage.

### Pha21 plays a positive role in virus resistance

To analyze whether Pha21 plays a role in virus resistance, we first analyzed the expression of *Pha21* in response to virus infection. We detected the RNA level of Pha21 in mock- or CymMV-inoculated *P. aphrodite* using qRT-PCR, and the results showed that the RNA of Pha21 is induced by CymMV infection (Fig. 3A). Furthermore, we transiently silenced or overexpressed Pha21 in CymMV-infected *P. aphrodite* to assay the effect on virus accumulation. Transient knockdown of Pha21 had no significant effect on CymMV accumulation (Fig. 3B); whereas transient overexpression of Pha21 decreased CymMV accumulation (Fig. 3C). In addition, we also generated transgenic *P. equestris* overexpressing Pha21 (35S::FLAG-Pha21). The expression of *Pha21* is higher in the two individual asexually propagated progeny derived from two T0 lines of transgenic *P. equestris* (Pha21#8 and Pha21#9) as compared to the non-transgenic lines (WT) (Fig. 3D). We further inoculated CymMV into individual progenies derived from two transgenic lines, and the result showed that the accumulation level of CymMV is decreased to 21% and 34% in the two transgenic lines (Pha21#8 and Pha21#9) as compared to the non-transgenic lines (WT) (Fig. 3D). Our data suggest that Pha21 plays a positive role in virus resistance.

**FIG 3.**
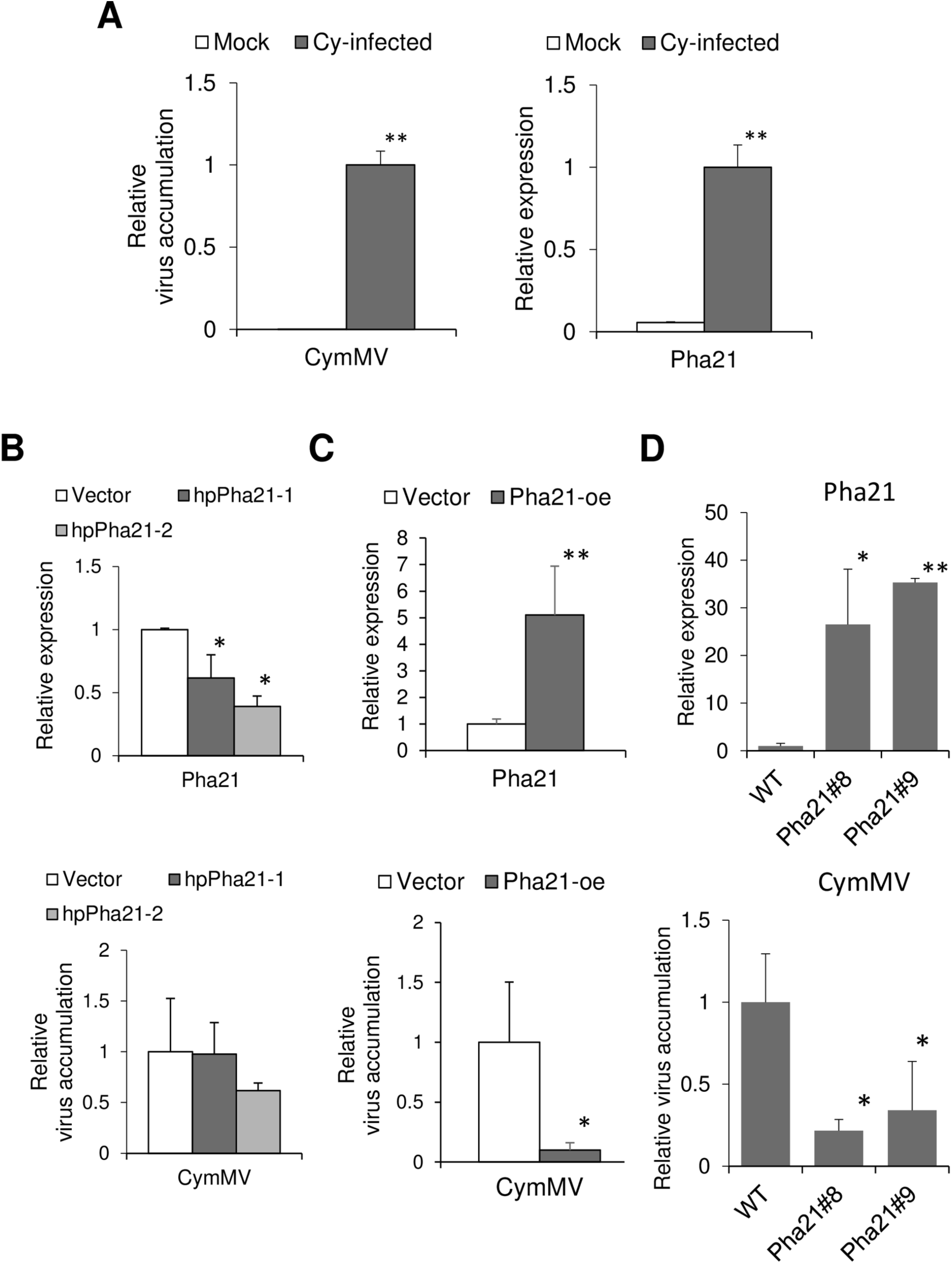
Pha21 is involved in virus accumulation. (A) qRT-PCR was performed to quantify the CymMV accumulation level and expression level of *Pha21* in healthy (Mock) and CymMV*-*infected (Cy-infected) *P. aphrodite.* The expression level in the CymMV*-*infected (Cy-infected) *P. aphrodite* was set to 1. Data represent mean ± SD; n = 3 biological replicates; **, *P* < 0.01; *, *P* < 0.05, Student’s t test compared to mock. (B-C) Transient silencing or overexpression of Pha21 in CymMV-infected plants. qRT-PCR was performed to quantify the expression level of *Pha21* and CymMV accumulation level from leaves of CymMV-infected *P. aphrodite* infiltrated with agrobacterium carrying vector (Vector); hairpin RNA (hpRNA) vector to knockdown Pha21 (hpPha21-1 and hpPha21-2; B); or overexpression vector of Pha21 (Pha21-oe; C). The expression level of plants infiltrated with control vector was set to 1. Data represent mean ± SD; n = 3 biological replicates; **, *P* < 0.01; *, *P* < 0.05, Student’s t test compared to Vector. (D) The wild-type (WT) *P. equestris* orchid and transgenic *P. equestris* (Pha21#8 and #9) were used for analysis. expression level of *Pha21* and CymMV were analyzed by qRT-PCR from leaves of WT or transgenic *P. equestris* (Pha21#8 and #9) inoculated with CymMV. The expression level of WT was set to 1. Data represent mean ± SD; n = 2 biological replicates; **, *P* < 0.01; *, *P* < 0.05, Student’s t-test compared to WT. (A-D) *PhaUbiquitin 10* was used as an internal control for normalization.

### Transgenic *Arabidopsis* overexpressing Pha21 enhances antiviral resistance

To understand whether Pha21-mediated antiviral immunity is conserved in plants, we generated the transgenic *Arabidopsis* (Col-0) overexpressing Pha21 (35S::FLAG-Pha21). Homozygous T3 plants derived from 3 T1 transgenic lines, At-Pha21#4, #5 and #6, were selected for further antiviral assay. The expression level of *Pha21* was confirmed by qRT-PCR on the homozygote progenies (Fig. 4A).

**FIG 4.**
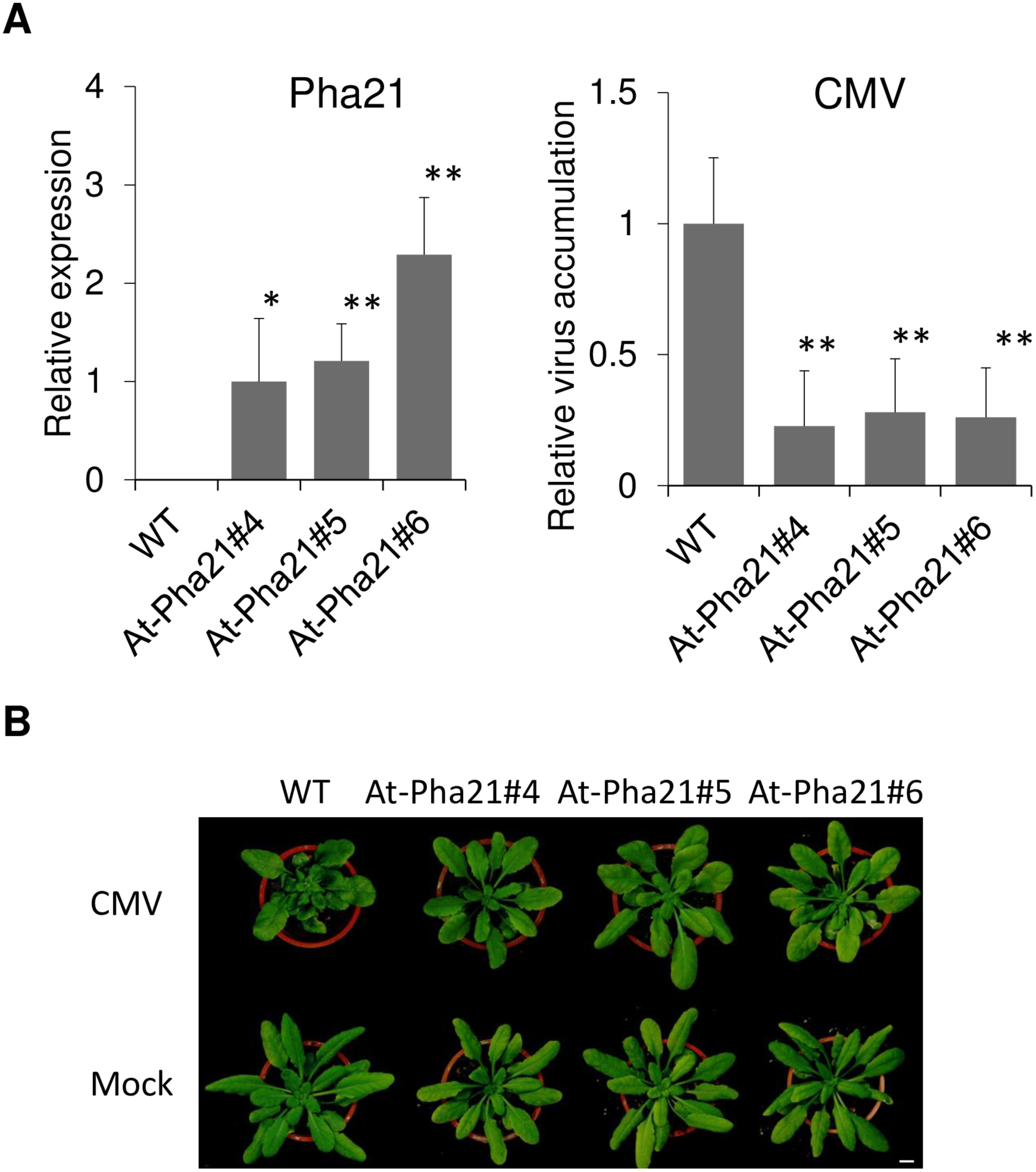
Overexpression of Pha21 in engineered transgenic *Arabidopsis* enhances resistance against viruses. (A) Expression level of *Pha21* and accumulation level of *Cucumber mosaic virus* (CMV) were analyzed by qRT-PCR from leaves of WT (Col-0) or transgenic *Arabidopsis* (At-Pha21#4, #5, and #6). Data represent mean ± SD; n = 5 biological replicates; **, *P* < 0.01; *, *P* < 0.05, Student’s t-test compared to WT. *AtActin* was used as an internal control for normalization. (B) Disease symptoms of WT (Col-0) or transgenic *Arabidopsis* (At-Pha21#4, #5, and #6) inoculated with CMV at 14 dpi. Images are one representative plant from five replicates; Scale bar, 1 cm.

We mechanically inoculated *Cucumber mosaic virus* (CMV) to wild-type (WT) and Pha21 overexpressing *Arabidopsis* (At-Pha21#4, #5 and #6). Fourteen days post-inoculation, the accumulation of CMV in WT and Pha21 overexpressing *Arabidopsis* were detected by qRT-PCR. The results showed that Pha21 overexpressing *Arabidopsis* decreased the accumulation of CMV (Fig. 4A) compared to the WT, and severe disease symptoms were observed on the WT but not in the Pha21 overexpressing *Arabidopsis* (Fig. 4B).

### Subcellular localization of Pha21

To better understand the role of Pha21 within cells, we first analyzed the subcellular localization of Pha21. We fused green fluorescent protein (GFP) in the N- or C-terminal of Pha21 driven by 35S promoter (pG-Pha21 and pPha21-G). The GFP-fused Pha21 was further transfected into protoplasts isolated from *P. aphrodite.* Twenty-four hours post-transfection, the localization of Pha21 was observed by using a confocal microscope. The results indicated that the C-terminal GFP fusion protein (Pha21-G) was observed in the nucleus in about 50% of cells; whereas the N-terminal GFP fusion protein (Pha21-G) showed a similar result to our GFP control vector, which had no nucleus-specific GFP (Fig. 5A). Our data suggested that Pha21 can move into the nucleus even without the predicated NLS signal.

**FIG 5.**
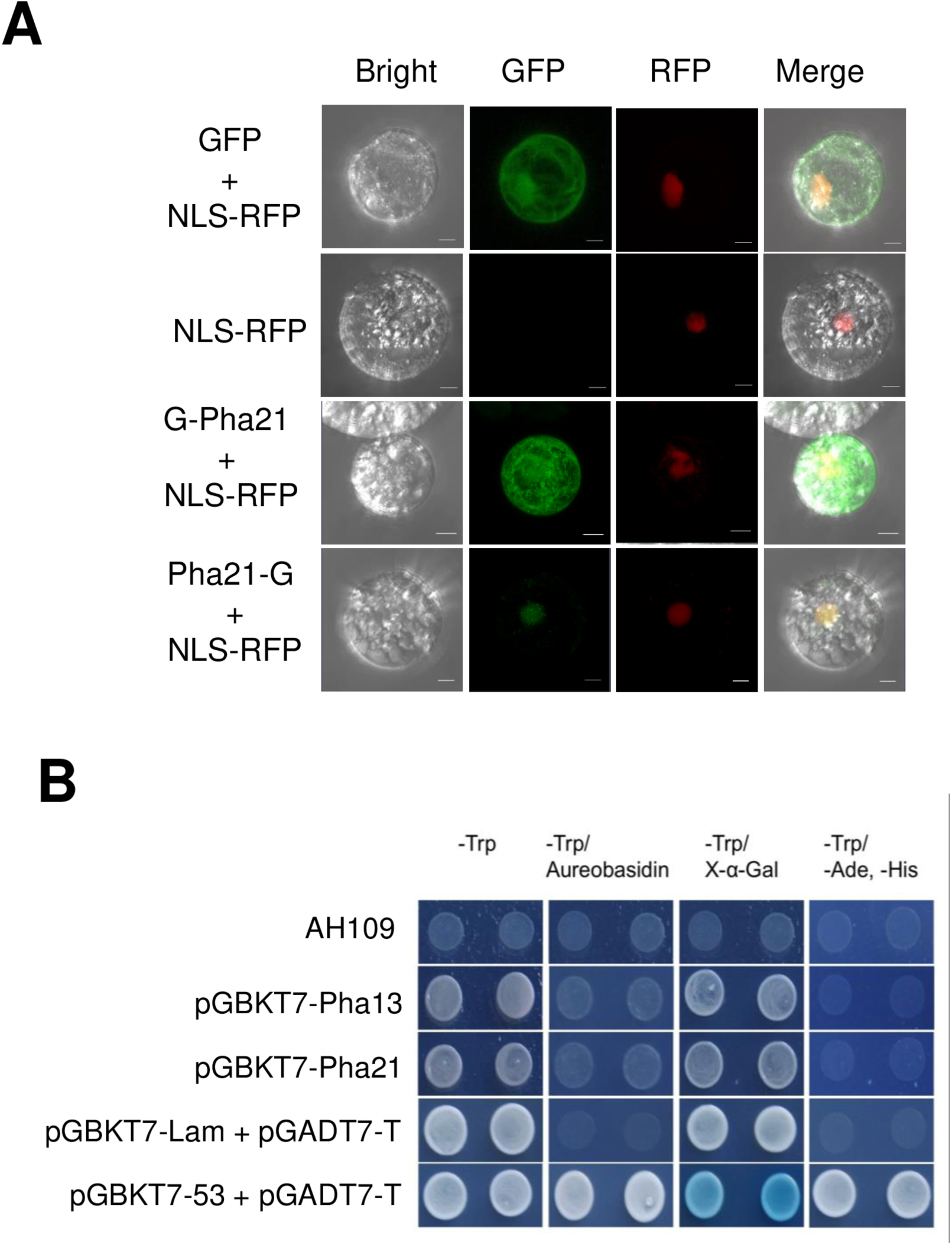
Subcellular localization of Pha21 and transcriptional activation analysis of Pha13 and Pha21. (A) Green fluorescent protein (GFP) or N- and C- terminal GFP-fused Pha21 (G-Pha21 and Pha21-G) were co-transfected with nucleus localization signal fused red fluorescence protein (NLS-RFP) into protoplasts of *P. aphrodite*. Fluorescence was detected by confocal microscopy after transfection. Scale bars represent 10 μm. (B) Yeast two-hybrid assay was used to analyze the transcriptional activation. AH109 is the yeast strain used in this assay. The pGBKT7-Pha13 and pGBKT7-Pha21 indicates that yeast encodes Pha13 or Pha21 fused to the Gal4 DNA binding domain. The pGBKT7-53, pGADT7-T, and pGBKT7-Lam provided in Matchmaker Gold Yeast Two-Hybrid System kit were used as a positive (pGBKT7-53 + pGADT7-T) and negative (pGBKT7-Lam + pGADT7-T) control. The -Trp indicates the medium without tryptophan. The -Trp/Aureobasidin was used to test for activation of the inositol phosphoryl ceramide synthase (AUR1-C) for Aureobasidin A resistance. The -Trp/X-α-Gal medium was used to test for activation of α-galactosidase, and -Trp/-Ade/-His indicate the selective medium without tryptophan, adenine, and histidine

### Pha21 and Pha13 does not confer transcriptional activation ability in yeast two-hybrid assay

Since both Pha21 and Pha13 have the ability to move into the nucleus (44), we tried to understand whether Pha21 and Pha13 function as transcription factors. Therefore, we analyzed its transcriptional activation ability through yeast two-hybrid (Y2H) assay. We fused Pha21 and Pha13 to the Gal4 DNA binding domain to generate pGBKT7-Pha21 and pGBKT7-Pha13. We transformed the pGBKT7-Pha21 or pGBKT7-Pha13 in the yeast strain AH109 without any Gal4 transcriptional activation domain (AD) and further analyzed the transcriptional activation ability through the analysis of the activation of reporter genes in yeast. As shown in Fig. 5B, yeast with pGBKT7-Pha21 or pGBKT7-Pha13 cannot grow on the -Trp/Aureobasidin and -Trp/-Ade/-His medium, and the colonies did not turn blue on X-α-Gal containing medium.

### Pha21 exhibits E3 ligase activity in which the A20 domain plays a major role

Several A20 and/or AN1 domain containing proteins confer E3 ligase activity (44–46, 53). To determine whether Pha21 functions as an E3 ligase, *in vitro* self-ubiquitination E3 ligase activity assay was performed using FLAG-Ubiquitin (FLAG-Ub), human E1 (hE1), human E2 (UBCH2, hE2), and purified His-tagged Pha21 (Pha21-His). The poly-ubiquitinated Pha21 can be detected in the presence of FLAG-Ub, hE1, and hE2 using anti-FLAG antibody to detect FLAG-Ubiquitin. The results revealed that Pha21 confers E3 ligase activity (Fig. 6A).

**FIG 6.**
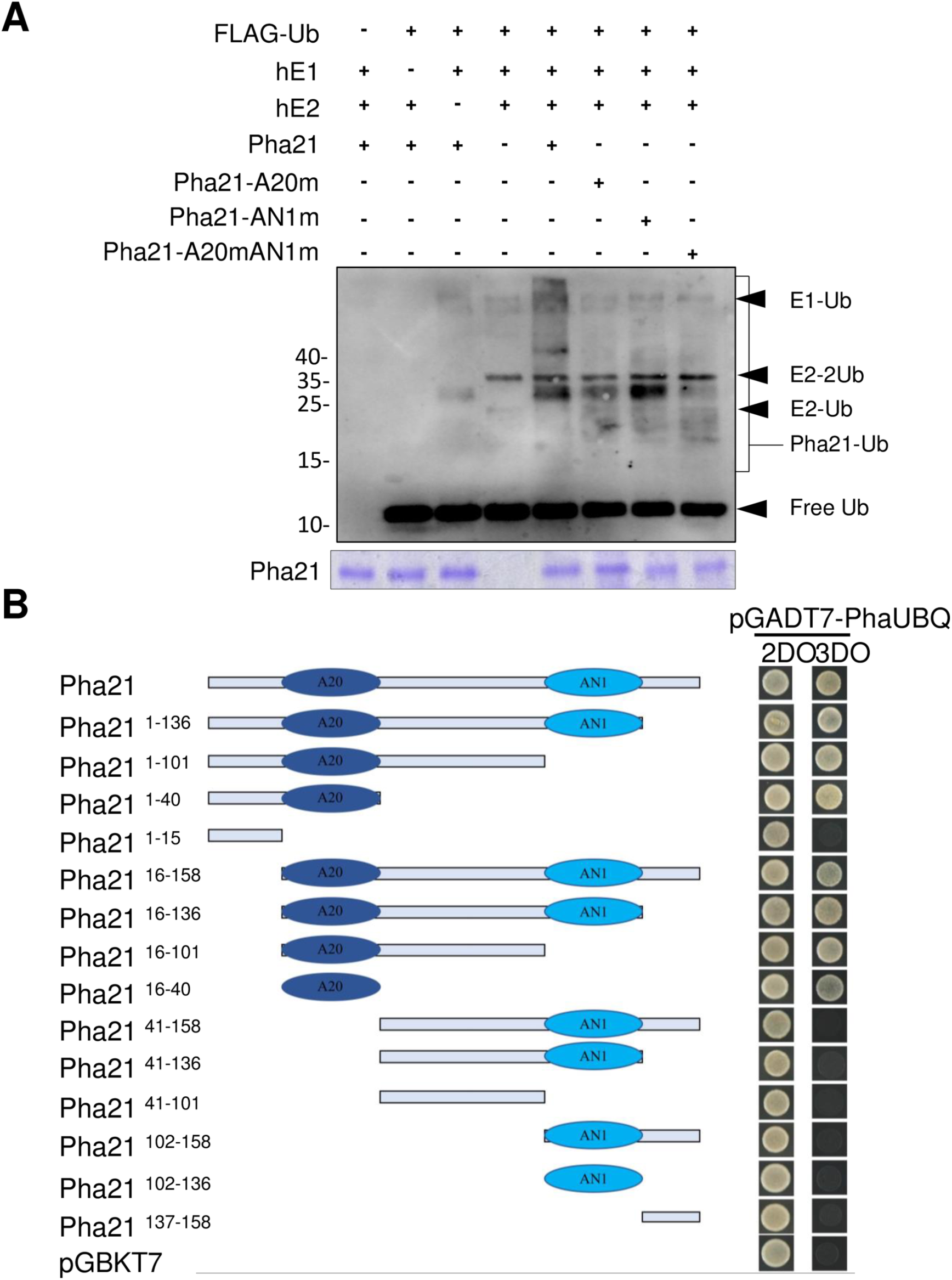
Pha21 confers self-ubiquitination E3 ligase activity and ubiquitin binding ability. (A) The *in vitro* self-ubiquitination E3 ligase activity assay was conducted on wild-type Pha21, A20 mutant (Pha21-A20m), AN1 mutant (Pha21-AN1m), A20/AN1 double mutant (Pha21-A20mAN1m) with or without FLAG-Ub, human ubiquitin-activating enzyme (hE1), or human ubiquitin-conjugating enzyme (hE2). Coomassie staining for Pha21 was used as a loading control. Ubiquitinated proteins were analyzed with immunoblotting using anti-FLAG antibodies. The ubiquitinated Pha21 are indicted as Pha21-Ub. E1 conjugated with one Ub (E1-Ub), E2 conjugated with one or two Ub (E2-Ub, E2-2Ub), and free Ub are indicated with a black arrow. (B) Yeast two-hybrid assay was performed to analyze the interaction between full-length and deletion mutant of Pha21 and ubiquitin. The schematic representations of full-length and deletion mutant of Pha21 are indicated. The open rectangle indicates the entire Pha21 protein. A20 domain and AN1 domain are indicated as a dark blue oval circle and a light blue oval circle, respectively. The full-length and deletion mutant of Pha21 cloned in pGBKT7 vector were co-transformed with pGADT7-PhaUBQ into yeast AH109 strain. The 2DO indicates the Leucine and Trptophan dropout selective medium and 3DO indicates the Leucine, Trptophan, and Histidine dropout selective medium. The pGBKT7 was used as a negative control.

Furthermore, we also analyzed the self-ubiquitination E3 ligase activity of the A20 and/or AN1 domain of Pha21. We substituted the conserved cysteine and histidine to glycine on the A20 and/or AN1 domains of Pha21-His (Fig 1B and C) to generate A20 domain mutant (Pha21-A20m), AN1 domain mutant (Pha21-AN1m), and the double mutant (Pha21-A20mAN1m) for E3 ligase activity analysis. As shown in Fig 6A, the major E3 ligase activity was conferred by the A20 domain. A20 domain mutant and double mutant of Pha21 showed lower self-ubiquitination E3 ligase activity than wild-type Pha21 and Pha21-AN1 mutant (Fig. 6A).

### A20 domain of Pha21 confers ubiquitin binding ability

A20 and/or AN1 proteins have also been shown to confer ubiquitin binding ability in animals and plants (44, 46, 47, 54, 55). To analyze whether Pha21 has ubiquitin binding ability, Y2H assay was performed to verify the interaction using Pha21 as bait and ubiquitin as prey. As shown in Fig 6B, Pha21 had a positive interaction with ubiquitin, suggesting that Pha21 conferred ubiquitin binding ability. In addition, to identify the ubiquitin binding domain of Pha21, a series of deletion mutants of Pha21 were generated, followed by Y2H assay. The result revealed that only truncated Pha21 fragments containing A20 domain (Pha21^1-136^, Pha21^1-101^, Pha21^1-40^, Pha21^16-158^, Pha21^16-136^, Pha21^16-101^, Pha21^16-40^) showed a positive interaction with ubiquitin, suggesting that the A20 domain of Pha21 is responsible for the ubiquitin binding ability (Fig. 6B).

### Interaction analysis between orchid SAPs

The A20/AN1 proteins may confer self-interaction and also interact with each other (56). Therefore, we also analyzed whether Pha21 and Pha13 confer self-interaction and also interact with each other. The Y2H assay was performed to verify the interaction between Pha21 and Pha13. Our Y2H results suggested that Pha21 and Pha13 did not interact with each other and no self-interaction of Pha21 or Pha13 was observed (Fig. 7).

**FIG 7.**
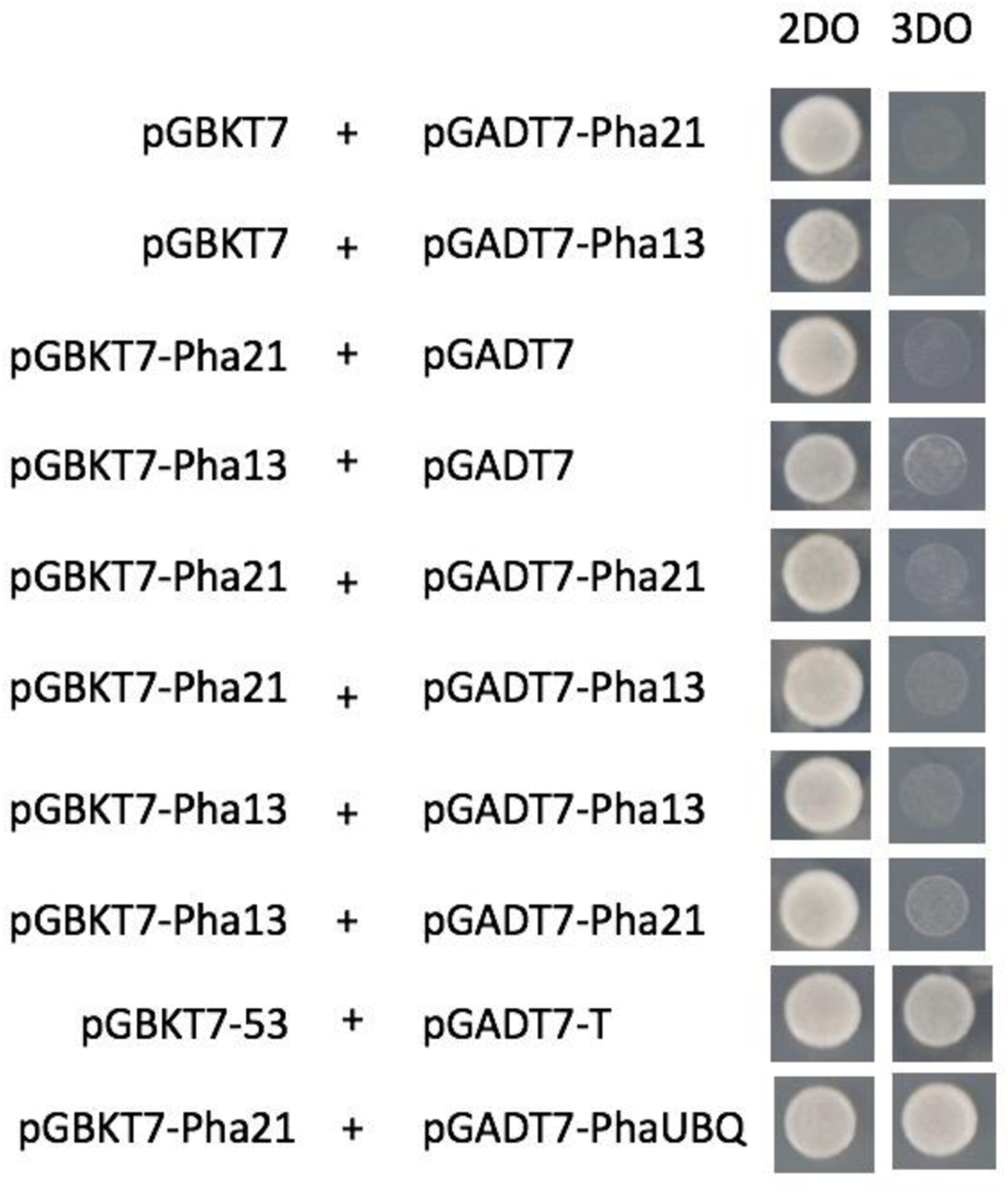
Interaction analysis of Pha21 and Pha13. Yeast two-hybrid assay was performed to analyze the interaction between Pha21 and Pha13. The pGBKT7 and pGADT7 were used as a negative control. The pGBKT7-53 and pGADT7-T (provided in the Matchmaker Gold Yeast Two-Hybrid System kit), and pGBKT7-Pha21 and pGADT7-PhaUBQ were used as a positive control. The 2DO indicates the dropout selective medium without Leucine and Tryptophan. The 3DO indicates dropout selective medium without the Leucine, Tryptophan, and Histidine.

### Pha21 is involved in the expression of SA-induced PhaNPR1-dependent and - independent genes

To further understand how Pha21 is involved in the antiviral immunity, we performed transient silencing (35S::hpPha21-2) and overexpression assay (35S::FLAG-Pha21) of Pha21 in *P. aphrodite*, and analyzed the expression of the previously identified PhaNPR1-dependent and -independent antiviral genes, *Phalaenopsis* homolog of *RNA dependent RNA polymerase 1* (*PhaRdR1)* and *Glutaredoxin C9 (PhaGRXC9)*, respectively (44). Our results indicated that transient silencing of Pha21 RNA increased the expression of *PhaRdR1*, while *PhaGRXC9* remained unchanged (Fig. 8A). Transient overexpression of Pha21 RNA decreased the expression of *PhaGRXC9*, but *PhaRdR1* remain unchanged (Fig. 8B). Our data suggest that Pha21 affects the expression of *PhaRdR1* and *PhaGRXC9*.

**FIG 8.**
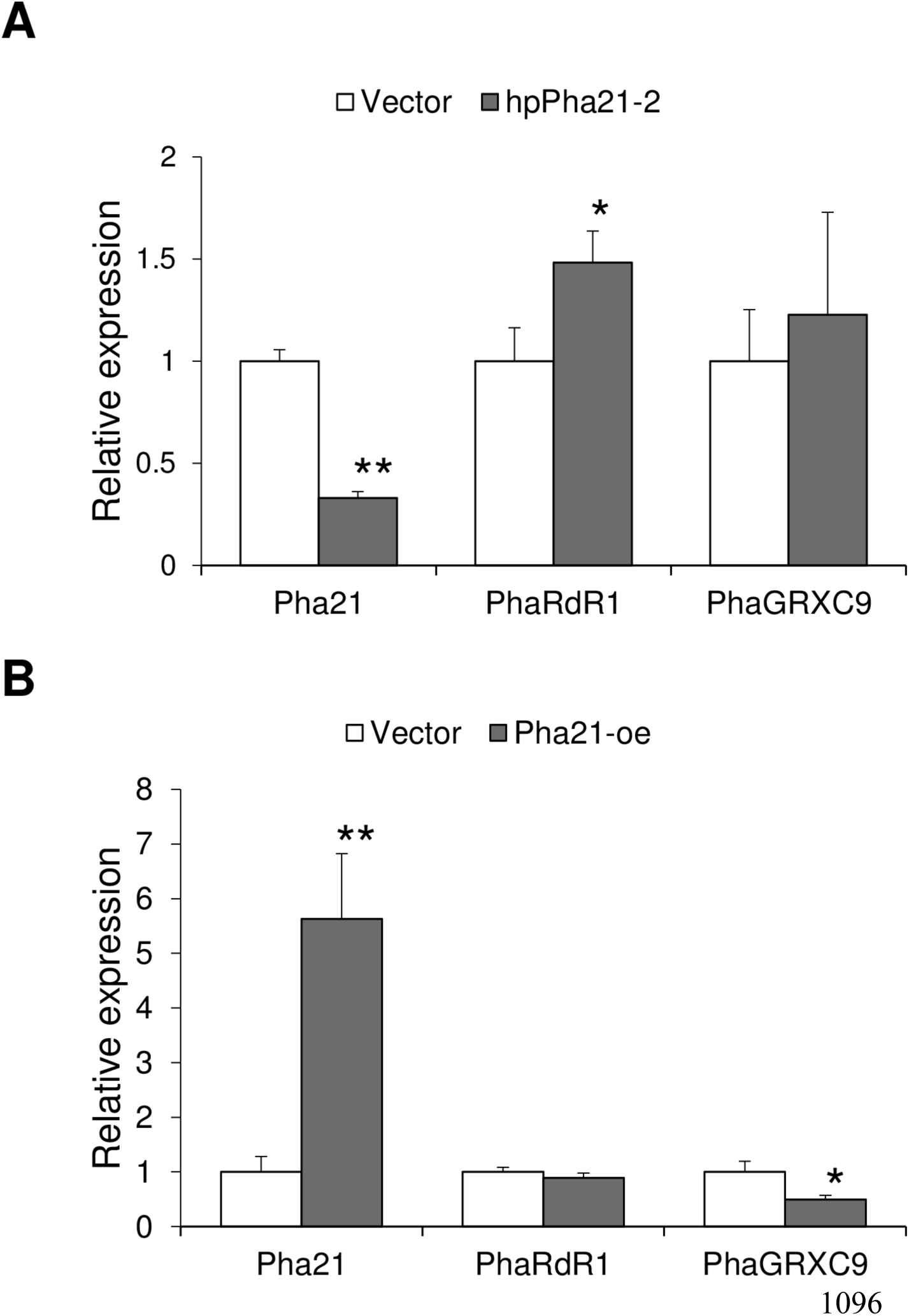
Pha21 is involved in the expression of PhaNPR1-dependent gene, *PhaRdR1,* and PhaNPR1-independent gene, *PhaGRXC9*. The expression levels of *Pha21, PhaRdR1*, and *PhaGRXC9* were analyzed by qRT-PCR from leaves of *P. aphrodite* infiltrated with agrobacterium carrying control vector (Vector) or hairpin RNA (hpRNA) vector to knockdown Pha21 (hpPha21-2; A), or to overexpress Pha21 (Pha21-oe; B). The RNA level of plants infiltrated with control vector was set to 1. Data represent mean ± SD; n = 3 biological replicates; **, *P* < 0.01; *, *P* < 0.05, Student’s t-test compared to Vector. (A-B) *PhaUbiquitin 10* was used as an internal control for normalization.

### Pha21 is involved in the expression of *PhaDCL4* and *PhaAGO1*

Because the RNAi pathway plays an important role in antiviral immunity (18, 19), we analyzed whether Pha21 is involved in the expression of core RNAi-related genes including the *Phalaenopsis* homolog genes of *RdR2* (*PhaRdR2*, PATC124544), *RdR6* (*PhaRdR6*, PATC131836), *DCL2* (*PhaDCL2*, PATC143544), *DCL4* (*PhaDCL4*, PATC150652), *AGO1* (*PhaAGO1*, PATC157237), and *AGO10* (*PhaAGO10*, PATC093469). Transient overexpression and silencing assay of Pha21 was performed in *P. aphrodite*. Overexpression of Pha21 increased the RNA expression of *PhaDCL4* and *PhaAGO1* (Fig. 9A); whereas transient silencing of Pha21 had no significant effect on the RNA expression of RNAi-related genes (Fig. 9B).

**FIG 9.**
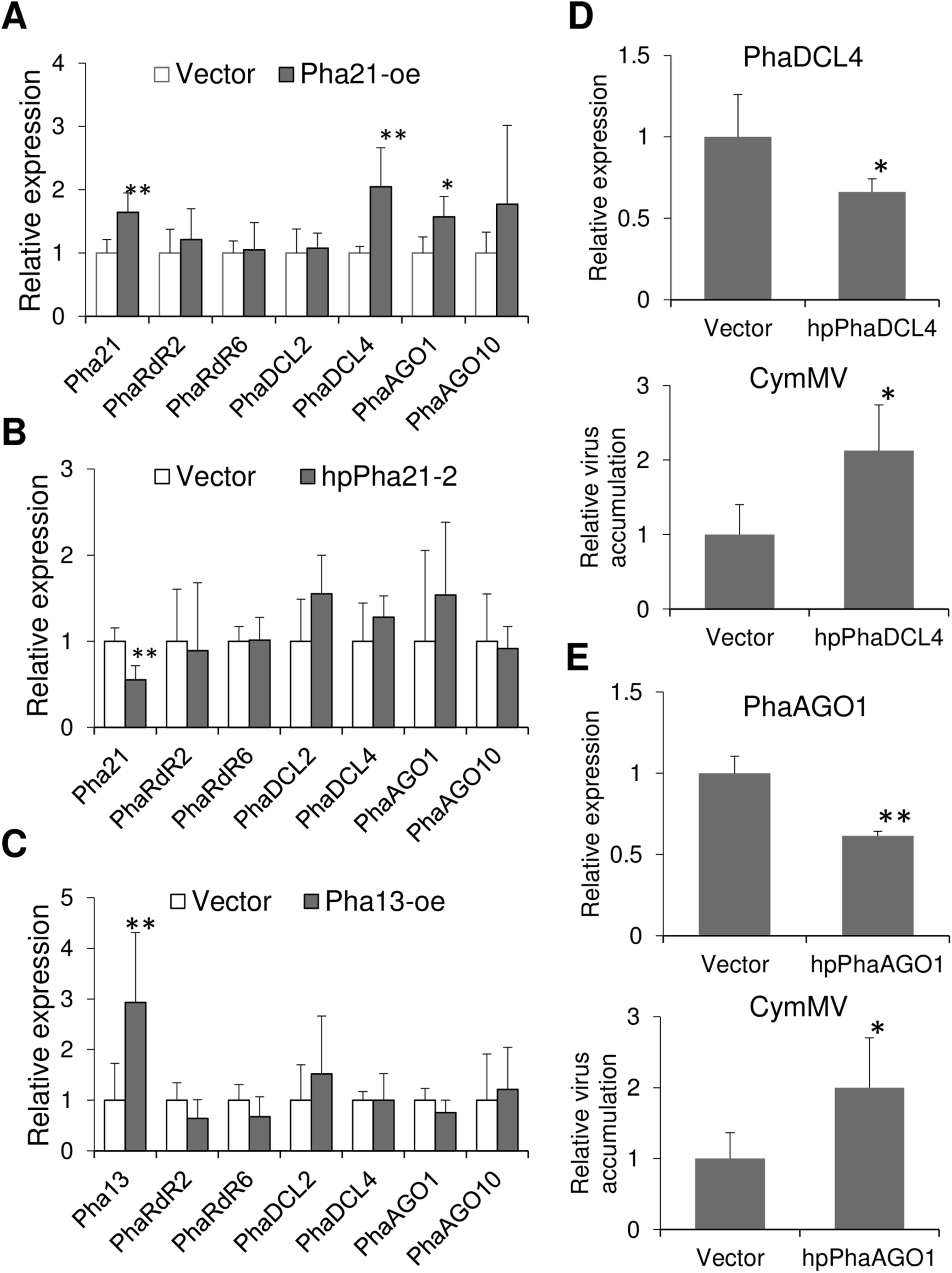
Pha21 is involved in the expression of *PhaDCL4* and *PhaAGO1*. (A-C) The expression levels of *Pha13*, *Pha21*, *PhaRdR2*, *PhaRdR6*, *PhaDCL2*, *PhaDCL4*, *PhaAGO1*, and *PhaAGO10* were analyzed by qRT-PCR from leaves of *P. aphrodite* infiltrated with agrobacterium carrying control vector (Vector); overexpression vector of Pha21 (Pha21-oe; A); hairpin RNA vector to knock down Pha21 (hpPha21-2; B); and overexpression vector of Pha13 (Pha13-oe; C). The expression level of plants infiltrated with control vector was set to 1. Data represent mean ± SD; n = 3 biological replicates; **, *P* < 0.01; *, *P* < 0.05, Student’s t-test compared to Vector. (D-E) The expression level of *PhaDCL4*, *PhaAGO1*, and CymMV accumulation level were analyzed by qRT-PCR from leaves of CymMV-infected *P. aphrodite* infiltrated with agrobacterium carrying vector (Vector); hairpin RNA (hpRNA) vector to knock down PhaDCL4 (hpPhaDCL4, D), or PhaAGO1 (hpPhaAGO1, E). The expression level of plants infiltrated with control vector was set to 1. Data represent mean ± SD; n = 3 biological replicates; **, *P* < 0.01; *, *P* < 0.05, Student’s t test compared to Vector. (A-E) *PhaUbiquitin 10* was used as an internal control for normalization.

For comparison, we also analyzed the effect of Pha13 on the RNAi-related genes. The results showed that transient overexpression of Pha13 has no significant effect on the expression of *PhaRdR2*, *PhaRdR6*, *PhaDCL2*, *PhaDCL4*, *PhaAGO1*, *and PhaAGO10* (Fig. 9C). Our results suggested that Pha21 is involved in the expression of *PhaDCL4* and *PhaAGO1*.

To see whether orchid DCL4 (PhaDCL4) and AGO1 (PhaAGO1) also play a role in the antiviral immunity, we transiently silenced PhaDCL4 and PhaAGO1 through delivering the hairpin RNA into *P. aphrodite* by agroinfiltration carrying hairpin RNA expression constructs, phpPhaDCL4 and phpPhaAGO1. Our results showed that transient silencing of PhaDCL4 and PhaAGO1 increased the accumulation of CymMV (Fig. 9D and E).

### The *Arabidopsis* homolog gene of *Pha21*, *AtSAP5*, is involved in the expression of *DCL4* and *AGO1*

Our previous phylogenic analysis revealed that both Pha21 and Pha13 are closely related to *Arabidopsis* AtSAP5 (accession number: AT3G12630) (44). Therefore, we also analyzed whether AtSAP5 is involved in the expression of *DCL4* and *AGO1*. We detected the expression of *DCL4* and *AGO1* in our previously generated transgenic *Arabidopsis* overexpressing AtSAP5 (AtSAP5-oe-4 and 11) and RNAi lines of AtSAP5 (AtSAP5-RNAi-3 and 7) by qRT-PCR (44). The expression level of *AtSAP5* was confirmed by qRT-PCR (Fig. 10A). Our results showed that slightly increased expression of *DCL4* and *AGO1* were observed in overexpression transgenic lines, AtSAP5-oe-11 and decreased expression of *DCL4* and *AGO1* were observed in the RNAi lines, AtSAP5-RNAi-3 and AtSAP5-RNAi-7 (Fig. 10B and C). Our results suggest that AtSAP5 is important in the expression of *DCL4* and *AGO1*.

**FIG 10.**
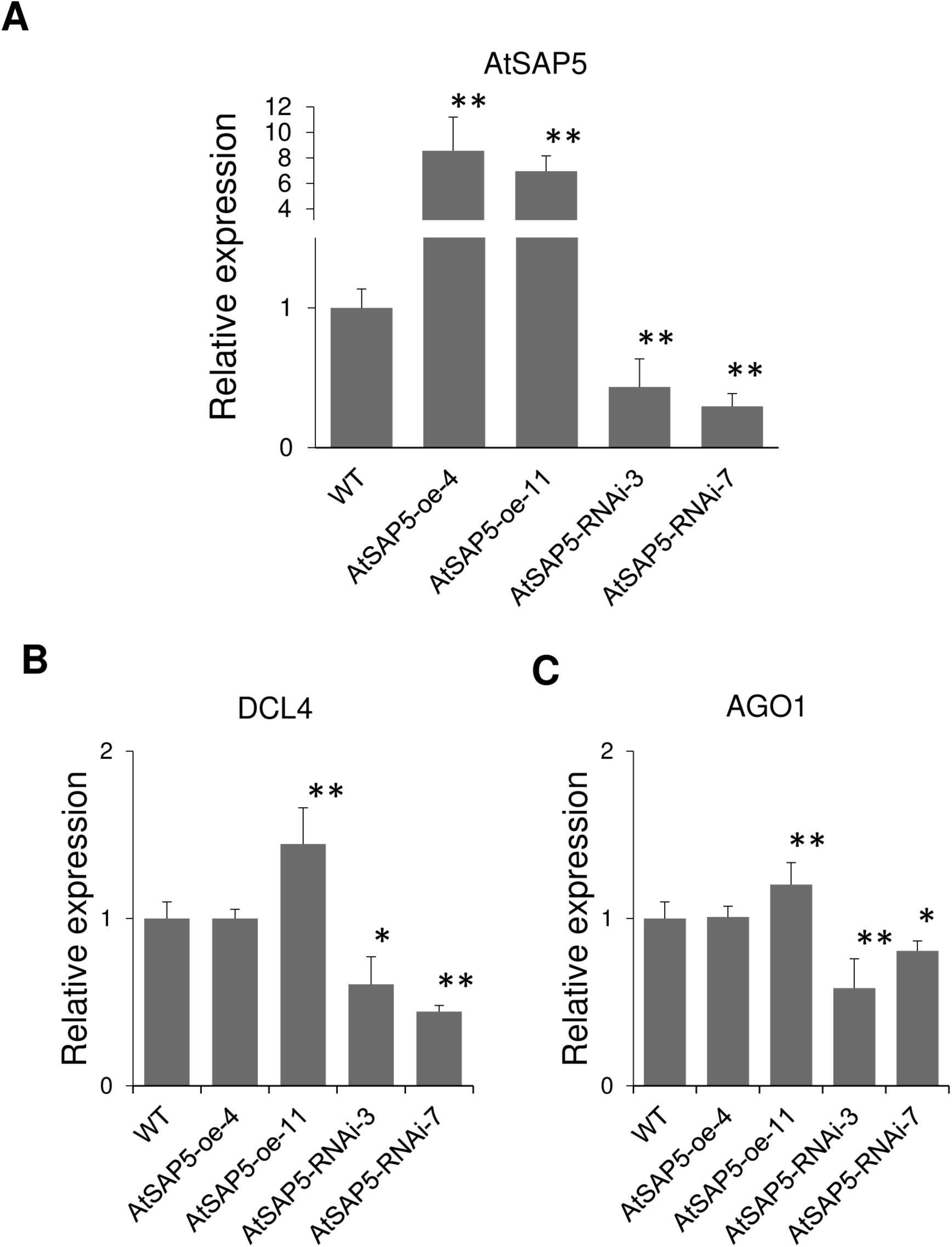
AtSAP5 is involved in the expression of *DCL4* and *AGO1*. (A-C) The expression levels of *AtSAP5, DCL4, AGO1* were analyzed by qRT-PCR in WT, AtSAP5 overexpression (AtSAP5-oe-4 and AtSAP5-oe-11) and RNAi lines (AtSAP5-RNAi-3 and AtSAP5-RNAi-7). Data represent mean ± SD; n = 5 biological replicates; **, *P* < 0.01; *, *P* < 0.05, Student’s t-test. Student’s t test compared to WT. *AtActin* was used as an internal control for normalization.

### A20 and AN1 domain of Pha21 play different roles in the expression of *PhaDCL4, PhaAGO1, PhaGRXC9* and virus resistance

Our previous results indicated that Pha21 is involved in the expression of *PhaDCL4*, *PhaAGO1*, and *PhaGRXC9* and virus accumulation (Figs 3C, 3D, 8B, and 9A). We further analyzed the roles of the A20 and/or AN1 domains of Pha21 in the expression of *PhaDCL4*, *PhaAGO1*, *PhaGRXC9* and virus accumulation by overexpression of Pha21 and Pha21 with mutation in the A20 and/or AN1 domain. Therefore, we generated Pha21 with A20 domain mutant (Pha21A20m), AN1 domain mutant (Pha21AN1m), and the double mutant (Pha21A20mAN1m) through the substitution of the conserved cysteine and histidine to glycine on A20 and/or AN1 domain (Fig. 1B and C). We transiently overexpressed wild-type Pha21 or the mutants (A20 and/or AN1 domain mutant) in healthy or CymMV pre-infected *P. aphrodite*, and then detected *PhaDCL4, PhaAGRO1, PhaGRXC9,* and CymMV. The results indicated that overexpression of Pha21 A20 mutant (Pha21A20m) but not AN1 mutant (Pha21AN1m) or A20/AN1 mutant (Pha21A20mAN1m) increased the expression of *PhaDCL4*, which is similar to the wild-type Pha21 (Fig. 11A). Overexpression of any Pha21 A20 and/or AN1 mutant showed similar results to wild-type Pha21 with respect to increased expression of *PhaAGO1* (Fig. 11A). Although overexpression of wild-type and any Pha21 A20 and/or AN1 mutant resulted in decreased expression of *PhaGRXC9* and CymMV accumulation compared to the vector control, overexpression of wild-type Pha21 showed a greater effect on the expression of *PhaGRXC9* and CymMV accumulation compared to any Pha21 A20 and/or AN1 mutant (Fig. 11A and B). Our data suggest that both the A20 and AN1 domains of Pha21 are required for expression of *PhaGRXC9* and the decrease of CymMV accumulation, and the AN1 domain plays a major role in the expression of *PhaDCL4*. They also suggest that neither domain is required for the expression of *PhaAGO1*.

**FIG 11.**
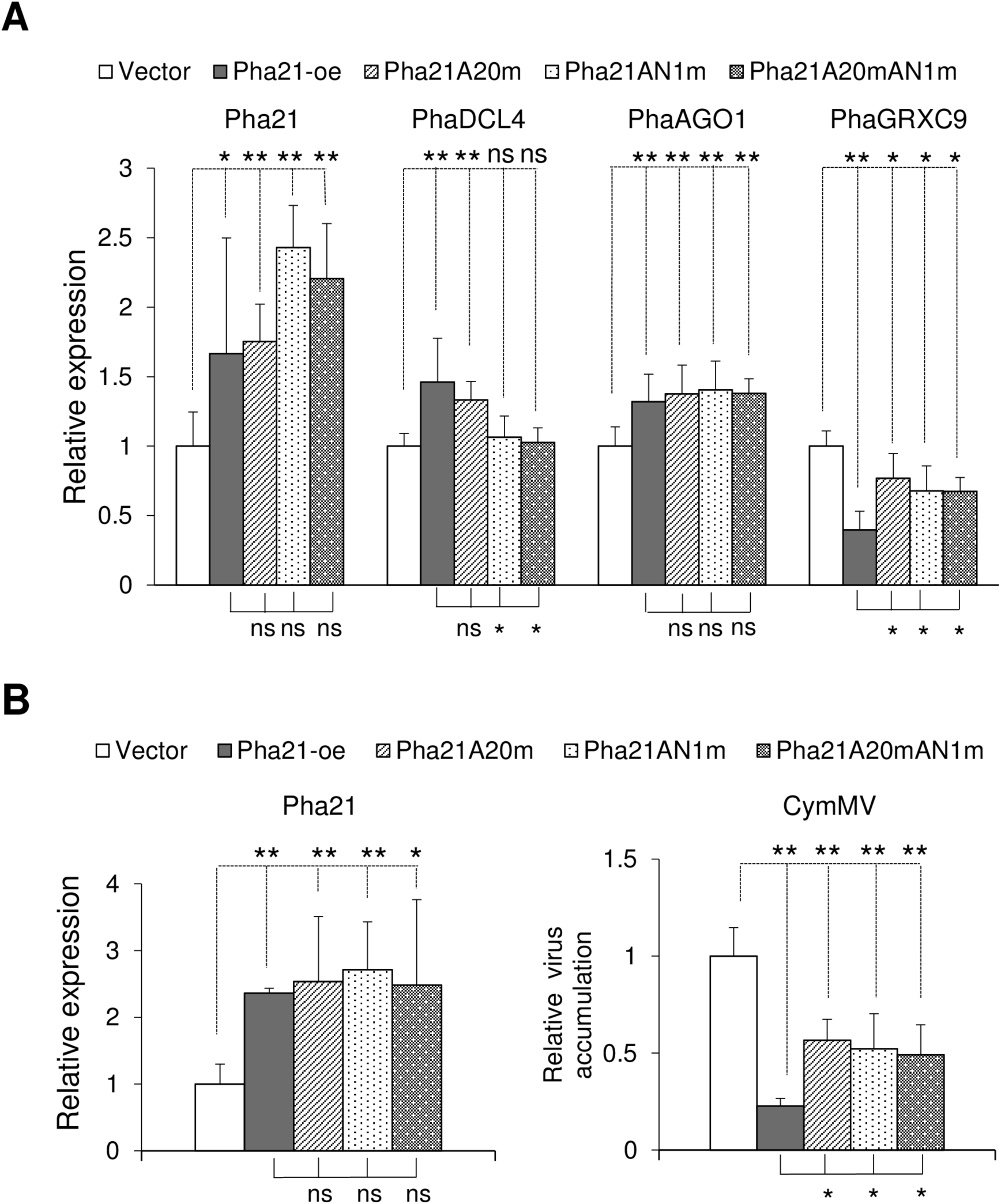
Pha21 A20 and AN1 domain play different roles in the expression of *PhaDCL4, PhaAGO1, PhaGRXC9* and virus resistance. The expression level of *Pha21*, *PhaDCL4, PhaAGO1*, *PhaGRXC9* and CymMV accumulation level were analyzed by qRT-PCR in leaves of healthy *P. aphrodite* (A), or CymMV pre-infected *P. aphrodite* (B) infiltrated with agrobacterium carrying control vector (Vector), overexpression clones of Pha21 (Pha21-oe), or the respective A20 and/or AN1 mutant clones (Pha21A20m, Pha21AN1m, and Pha21A20mAN1m). The RNA level of plants infiltrated with control vector was set to 1. Data represent mean ± SD; n = 3 biological replicates; **, *P* < 0.01; *, *P* < 0.05; ns, no significant difference, Student’s t test compared to Vector. (A-B) *PhaUbiquitin 10* was used as an internal control for normalization.

## DISCUSSION

### Plant A20/AN1 proteins, Pha13, Pha21 and AtSAP5 share similar biochemical properties and all involved in SA-mediated antiviral immunity

In this report, we performed detailed analysis of a previously identified orchid gene, *Pha21*, involved in SA and RNAi-mediated antiviral immunity. Pha21 shares high protein sequence identity to our previously identified Pha13 (Fig. S7 in Chang et al, 2018), and exhibits similar biochemical properties including self-E3 ligase, ubiquitin binding abilities (Fig. 6) and subcellular localization (Fig. 5A). Similar to Pha13, our data suggests that Pha21 is involved in the SA-signaling pathway downstream of SA and upstream of the expression of *PhaNPR1* (Fig. 2A, C and D). In addition, Pha13 and Pha21 also positively mediate antiviral immunity (44). Transgenic *Arabidopsis* overexpressing Pha21 also enhanced resistance to virus infection (Fig 4) (44). In addition, our previous report and the data presented here indicate that *Arabidopsis* AtSAP5 is also induced by SA at the early stage, and is involved in expression of similar sets of SA mediated immune responsive genes, and positively regulates antiviral immunity (44).

### Pha13-mediated expression of *PhaGRXC9* and *PhaRdR1* are dominant during SA treatment

Our previous study indicated that Pha13 and AtSAP5 plays a positive role in the expression of genes involved in the RNAi pathway and SA-governed oxidoreductases, *RdR1* and *GRXC9*, respectively (44). We demonstrated that both PhaRdR1 and PhaGRXC9 play positive roles in antiviral immunity (44). In this study, we found that Pha21 has an opposite role to Pha13 in the expression of *PhaRdR1* and *PhaGRXC9* (Fig. 8). As our data indicates that the RNA level of Pha13 is approximately 15 times higher than Pha21 in leaves of orchids as measured by digital PCR (Fig. 1E), we suggest that the effect of Pha21 on the expression of *PhaRdR1* and *PhaGRXC9* is minor, and Pha13-mediated positive expression of *PhaGRXC9* and *PhaRdR1* are dominant during SA treatment.

### Pha21 and AtSAP5 are all involved in expression of key genes in RNAi pathway, but regulation of *PhaDCL4* and *PhaAGO1* by Pha21 in orchid is different from regulation of *DCL4* and *AGO1* by AtSAP5 in *Arabidopsis*

Our data indicates that similar to *Arabidopsis* DCL4 and AGO1, orchid PhaDCL4 and PhaAGO1 also play a similar positive role in antiviral immunity in orchids as a slight decrease in the RNA of PhaDCL4 or PhaAGO1 (approximately 40% decrease) has a prominent effect on CymMV accumulation (approximately 100% increase) (Fig. 9D and E), (19). Interestingly, our data suggests that Pha21 regulates antiviral immunity partly through the effect on the RNAi pathway, as transient overexpression of Pha21 (1.6-folds) but not Pha13 increased the expression of two *Phalaenopsis* orchid homolog genes of *DCL4* (*PhaDCL4*, 2-fold) and *AGO1* (*PhaAGO1*, 1.6-fold) (Fig. 9A and C).

Although the effect of AtSAP5 on expression of *DCL4* and *AGO1* is not as prominent as Pha21, our data shown here also indicates that AtSAP5 is involved in the expression of *DCL4* and *AGO*1. In both of our silencing lines, AtSAP5-RNAi-3 and −7 showed noticeable effect on the expression of *DCL4* and *AGO1* (Fig. 10), but that only one overexpression line, AtSAP5-oe-11 (expression increased about 7 times over AtSAP5) slightly increased the expression of *DCL4* (40% increase) and *AGO1* (20% increase). This suggests that AtSAP5 is still important for the maintenance of *DCL4* and *AGO1* expression.

Collectively, our data suggests that although Pha13 and Pha21 both participate in SA regulated immunity, they act differently in triggering downstream immune responsive genes.

### Pha13 and Pha21 showed no transcriptional activation activity by Y2H assay

Although Pha21 and Pha13 are involved in expression of *PhaGRXC9*, *PhaRdR1*, *PhaDCL4* and *PhaAGO1*, our Y2H analysis showed that Pha21 and Pha13 have no transcriptional activation activity (Fig. 5B).

### A20 confers most E3 ligase and ubiquitin binding activity but the AN1 domain also plays important roles in expression of immune responsive genes

Similar to other analyzed A20/AN1 proteins, our biochemical assay indicated that both Pha21 and Pha13 confer E3 ligase activity and ubiquitin binding ability, and A20 domain of Pha21 and Pha13 is the major domain exhibiting self-E3 ligase and ubiquitin binding activity (Fig. 6) (44). It has been well demonstrated that human A20 proteins (without an AN1 domain) regulate the master immune transcription factor NF-kB through ubiquitin editing activity including E3 ligase and ubiquitin binding ability on different substrates (53). It is likely that Pha21 and Pha13 may function in a similar manner to human A20 indirectly, rather than directly function as transcription factors. Our previous domain functional analysis revealed that both the A20 and AN1 domains of Pha13 are required for expression of *PhaRdR1*, *PhaGRXC9*, and virus accumulation, and the AN1 domain of Pha13 is involved in the expression of *PhaNPR1* (44). Here we also showed that A20 and AN1 domains of Pha21 are both required for expression of *PhaGRXC9* and CymMV accumulation, and the AN1 domain plays a more important role in the expression of *PhaDCL4* (Fig. 11). This indicates that although A20 confers most E3 ligase and ubiquitin binding activity, the AN1 domain also plays important roles in expression of immune responsive genes. Interestingly, our data suggests that neither domain of Pha21 is required for the expression of *PhaAGO1*, which suggests that protein regions other than A20 and AN1 domain in Pha21 are also involved in the expression of *PhaAGO1* (Fig. 11A).

### A20/AN1 proteins serve as important modulators in plant antiviral immunity

Our previous data and the findings presented here indicate that A20/AN1 protein-mediated antiviral immunity is conserved among plants, and A20/AN1 proteins may work alone (AtSAP5) or in a cooperative manner (Pha13 and Pha21) in antiviral immunity to induce the expression of the SA-mediated immune responsive genes, including *NPR1*, NPR1-independent oxidoreductases gene (*GRXC9*), and genes involved in the RNAi pathway *(RdR1*, *AGO1* and *DCL4*) (Fig. 12).

**FIG 12.**
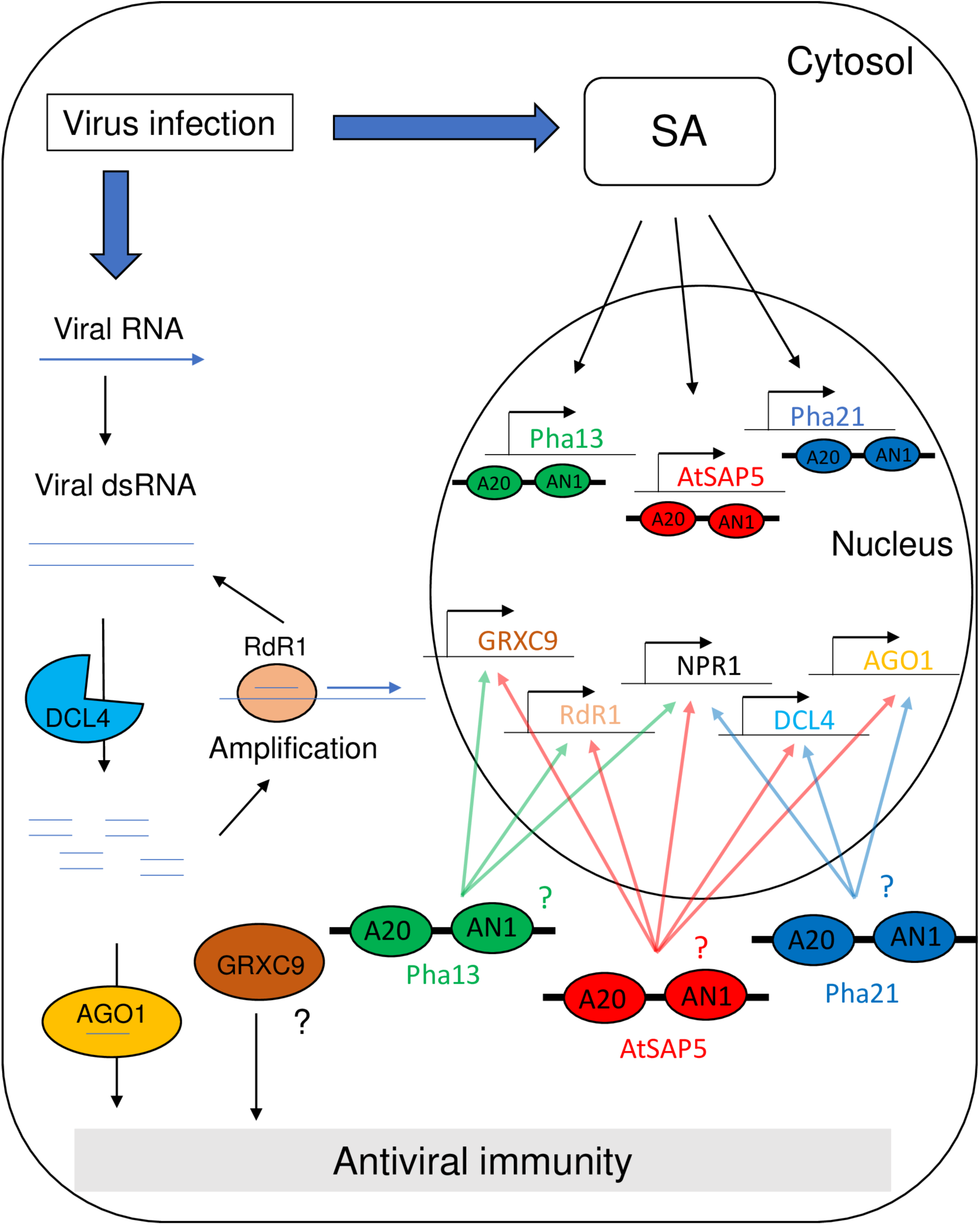
The role of A20/AN1 proteins in antiviral immunity. A model illustrating the roles of SA-induced-A20/AN1 proteins Pha13 and Pha21 from *Phalaenopsis aphrodite,* and AtSAP5 from *Arabidopsis thaliana* in antiviral immunity. Virus infection caused accumulation of SA and led to the activation of *Pha13*, *Pha21*, and *AtSAP5*, allowing it to trigger the SA-and RNAi-mediated immune responsive genes including *NPR1*, *GRXC9*, *DCL4*, *AGO1*, and *RdR1* in antiviral immunity.

## MATERIALS AND METHODS

### Plant materials and growth conditions

Orchid variety, *Phalaenopsis aphrodite var. Formosa*, was purchased from Taiwan Sugar Research Institute (Tainan, Taiwan). All orchid plants we used including

*P. aphrodite, P. equestris* and transgenic *P. equestris* (35S::FLAG-Pha21) were first analyzed for the infection with two prevalent orchid viruses, *Odontoglossum ringspot virus* (ORSV) and CymMV, as detected by RT-PCR with primer pairs, ORSV-F/ORSV-R and CymMV-F/CymMV-R (Table 1). Plants free from ORSV and CymMV were maintained in greenhouse conditions with a controlled 12-h photoperiod (200 μmol m^-2^s^-2^) at 25°C/25°C (day/night). The wild-type (WT) *Arabidopsis* WT (Col-0) and all transgenic *Arabidopsis* were maintained in a greenhouse with a controlled 12-h photoperiod (200 μmol m-2s-2) at 22°C/22°C (day/night) for three to four weeks before analysis. *Cucumber mosaic virus* (CMV) isolate 20 was maintained in the *Arabidopsis* (Col-0) as inoculation source in our study.

**Table 1.**
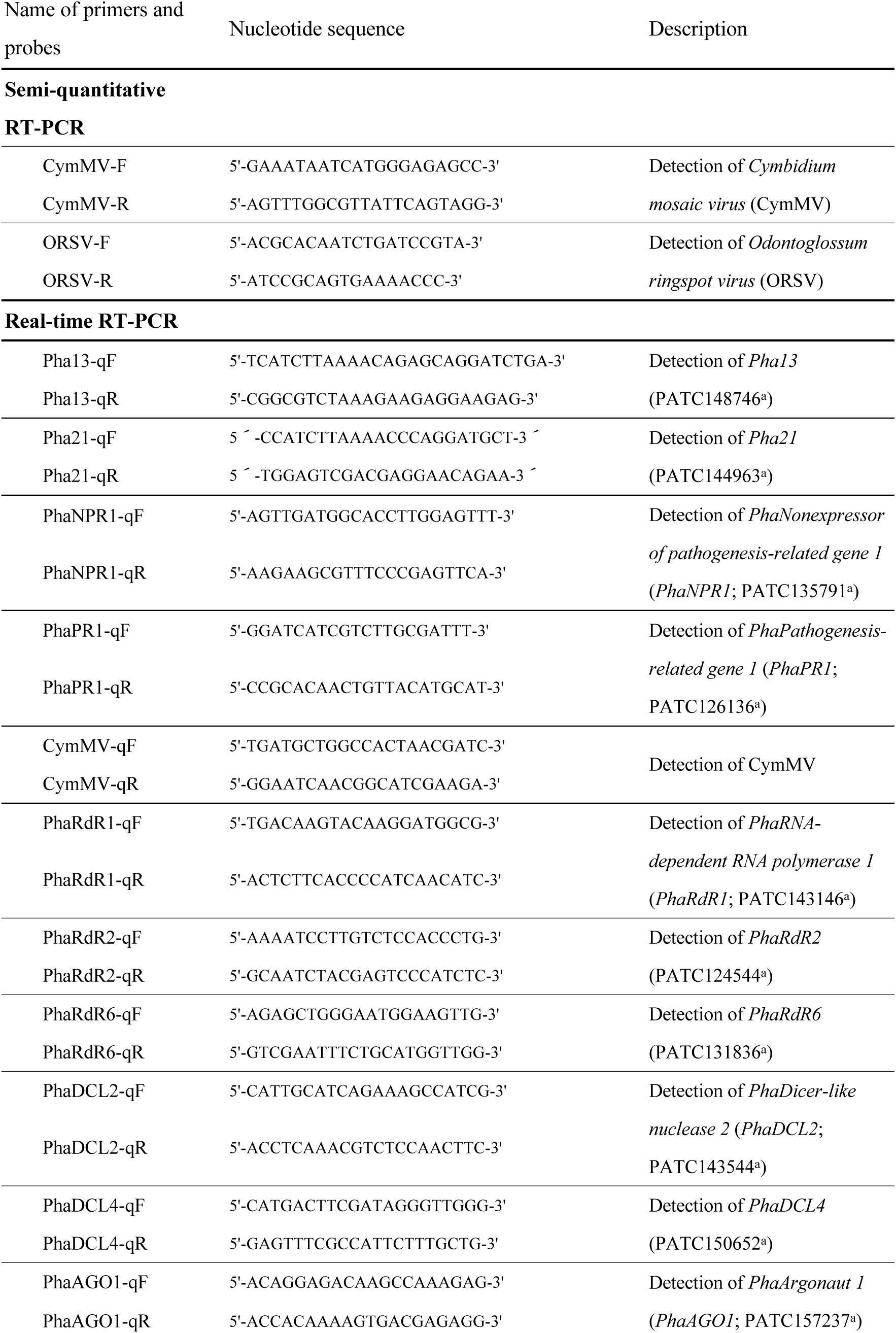
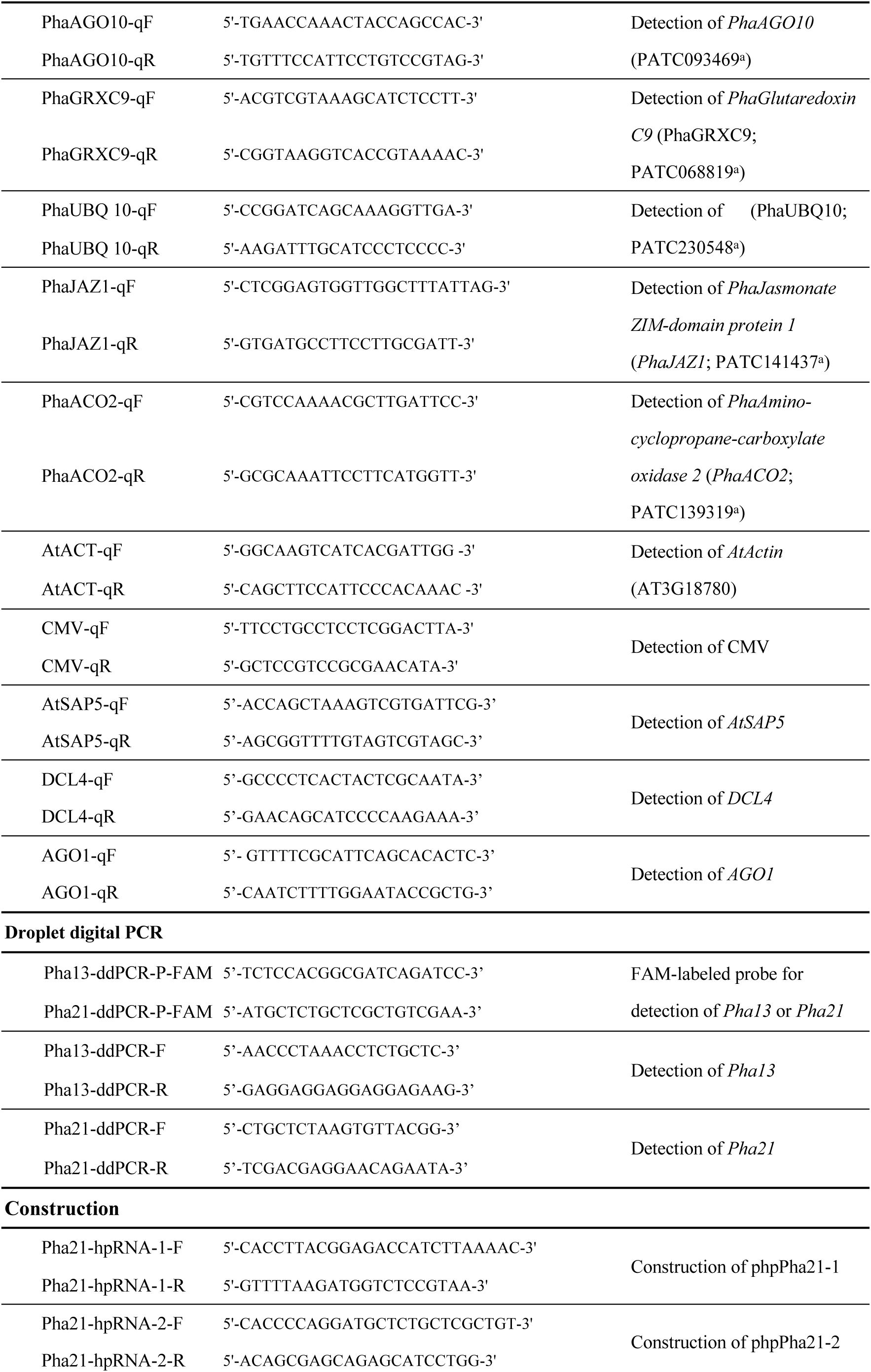
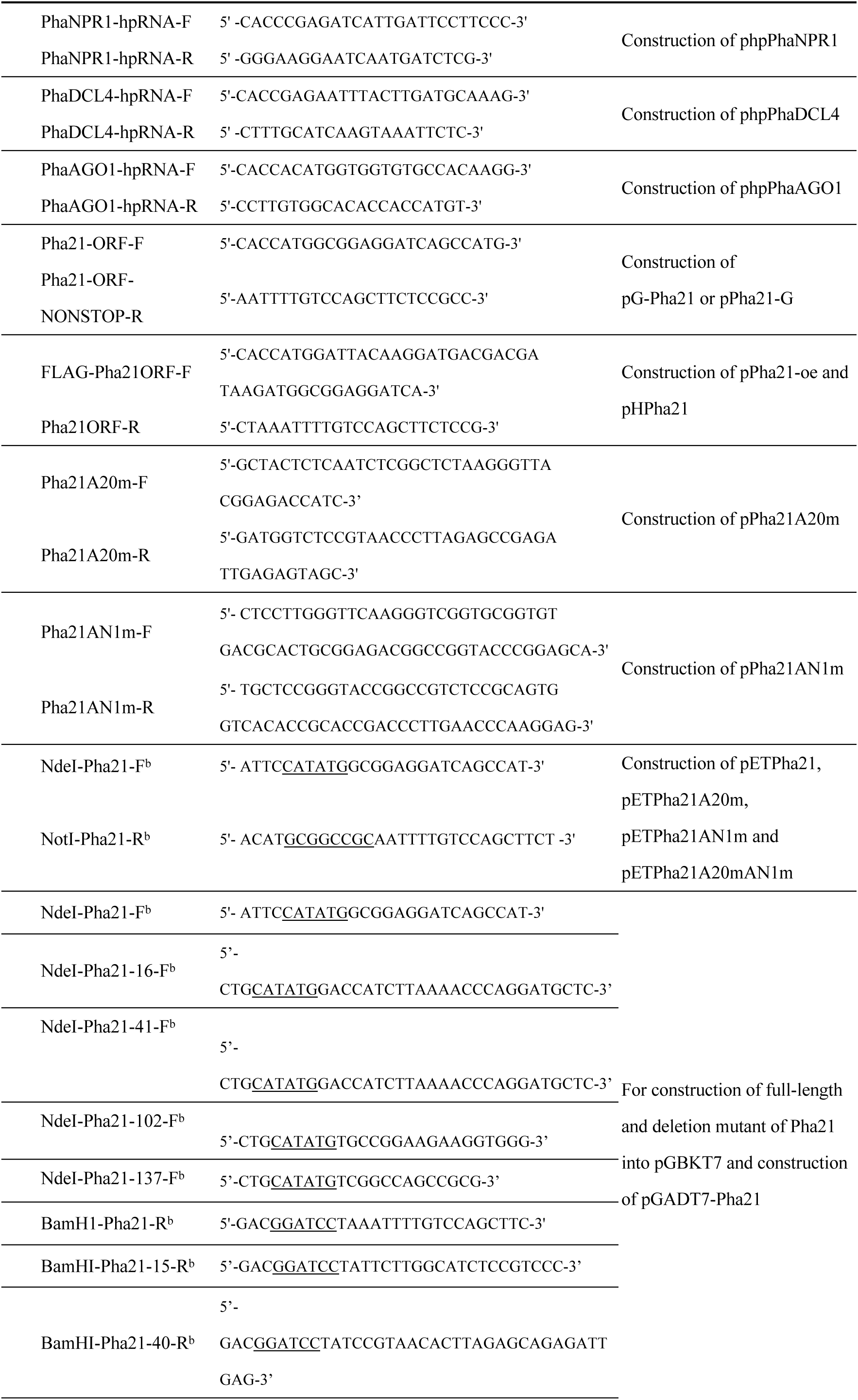
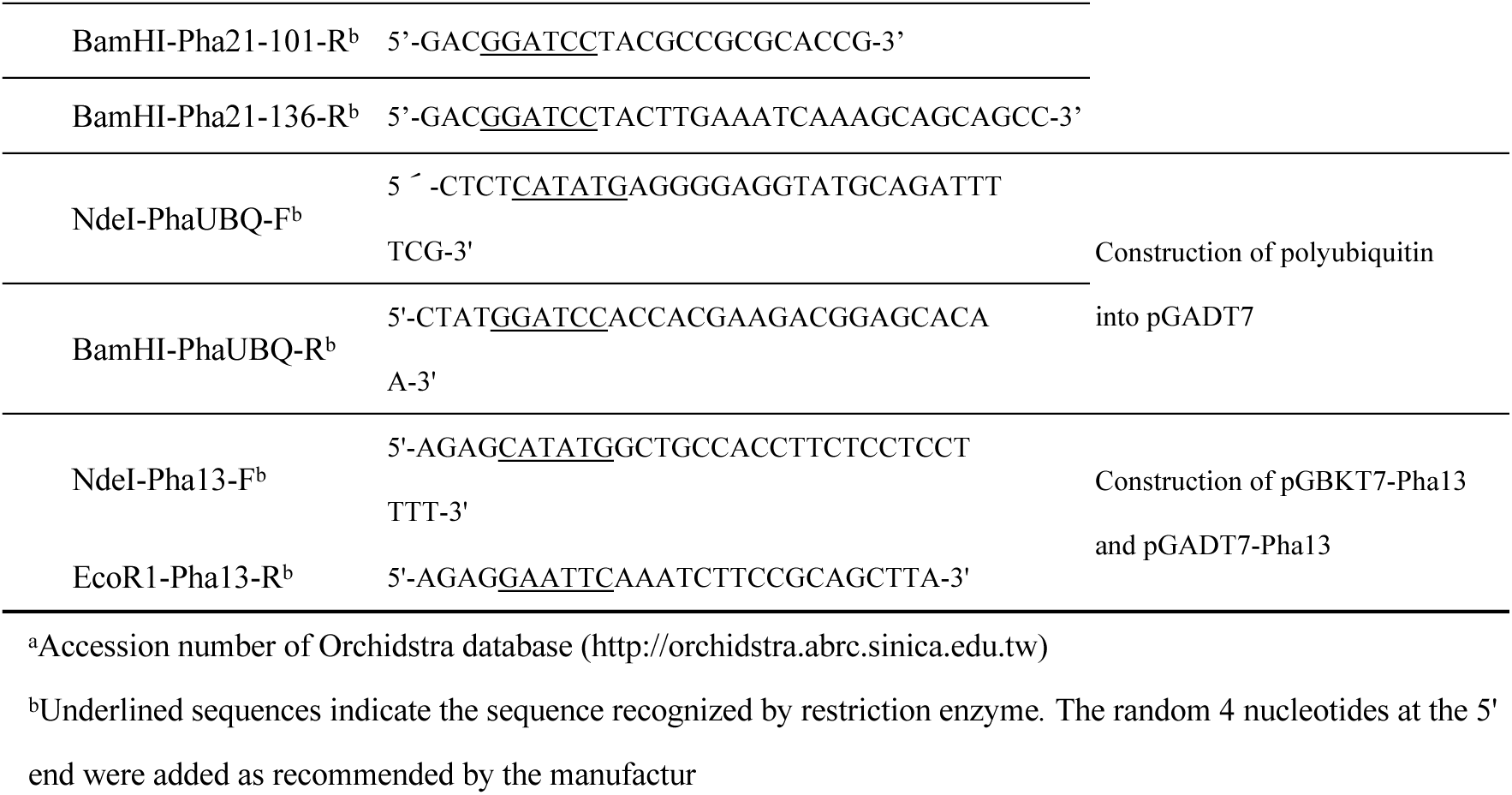
Primers and probes used in this study

### Sequence analysis

The amino acid sequence of Pha21 was analyzed by the PROSITE database, ExPASy Proteomics Server (http://ca.expasy.org/), and Conserved Domain Database of NCBI database (http://www.ncbi.nlm.nih.gov/).

### Phytohormone treatment

Sodium salicylate (50 mM) (Sigma, St. Louis, MO, USA), methyl jasmonate (45 μM) (Sigma), and aminocyclopropanecarboxylic acid (660 μM) (Sigma) were directly rubbed on leaves of *P. aphrodite* by cotton swab. Leaf samples were collected at 0 h, 1 h, 3 h, 6 h, 12 h, 24 h, 48 h and 72 h after treatment.

### RNA isolation and real-time quantitative RT-PCR (qRT-PCR) detection

Total RNA was extracted using the TOOLSmart RNA Extractor (BIOTOOLS, Taiwan) as described previously (44). The cDNA template for qPCR was synthesized from 500 ng of DNA-free RNA and oligo (dT) using PrimeScript RT Reagent Kit (Perfect Real Time), following the manufacturer’s instructions (Takara Bio, Shiga, Japan). qPCR was performed using SYBR Green PCR Master Mix (Applied Biosystems, Foster, CA, USA)) with ABI Prism 7500 sequence detection system (Applied Biosystems). Target gene PCR products were sequenced to validate the correct analysis of gene targets. *PhaUbiquitin 10* or *AtActin* was used as an internal quantification control. Quantification of target gene expression was calculated according to the manufacturer’s instructions (Applied Biosystems). The primer pairs used in this study are listed in Table 1.

### Droplet digital PCR

Total RNA was extracted using the TOOLSmart RNA Extractor (BIOTOOLS, Taiwan) as described previously (44). For droplet digital PCR (ddPCR), DNA-free RNA (1 μg) and oligodT primer were used for cDNA synthesis with PrimeScript RT Reagent Kit (Perfect Real Time) following the manufacturer’s instructions (Takara Bio). An amount of 5 μl of 25X dilute cDNA was used as a template for ddPCR reaction (total 20 μl) following the manufacturer’s instructions (Bio-Rad, Hercules, CA, USA). The sequence of FAM-labeled Taqman probes and primer pairs of Pha13 and Pha21 for ddPCR are listed in Table 1.

### Construction of transient silencing vector

For construction of transient silencing vector of Pha21, the oligonucleotide pairs Pha21-hpRNA-1-F/Pha21-hpRNA-1-R and Pha21-hpRNA-2-F/Pha21-hpRNA-2-R (Table 1) were used to amplify the hairpin dsDNA fragments. The hairpin dsDNA fragments were cloned into the Gateway entry vector pENTR/D-TOPO (Thermo Fisher-Scientific, Waltham, MA, USA) following the manufacturer’s instructions to generate pENTR-Pha21-hpRNA-1 and pENTR-Pha21-hpRNA-2. Then, LR Gateway cloning reaction (Thermo Fisher-Scientific) was conducted to transfer the hairpin RNA fragments from pENTR-Pha21-hpRNA into 35S promoter driven pB7GWIWG2(I) (57) to obtain phpPha21-1 and phpPha21-2. The method for construction of the transient silencing vector of *PhaNPR1, PhaDCL4, and PhaAGO1* (phpPhaNPR1, phpPhaDCL4, and phpPhaAGO1) were similar to that described above, except the oligonucleotide pairs PhaNPR1-hpRNA-F/PhaNPR1-hpRNA-R, PhaDCL4-hpRNA-F/PhaDCL4-hpRNA-R, and PhaAGO1-hpRNA-F/ PhaAGO1-hpRNA-R (Table 1) were used to generate the hairpin dsDNA fragments.

### Construction of transient overexpression vector of Pha21, Pha21 A20 and/or AN1 domain mutant

To construct Pha21 transient overexpression vector, total *P. aphrodite* RNA was used as a template to amplify the N-terminus FLAG-tagged Pha21 by RT-PCR with the primer pairs FLAG-Pha21ORF-F/Pha21ORF-R (Table 1). The amplified fragment was cloned into Gateway entry vector pENTR/D-TOPO (Invitrogen) to generate pENTR-FLAG-Pha21 following the manufacturer’s protocol. Subsequently, LR Gateway cloning reaction (Invitrogen) was performed to transfer the FLAG-Pha21 fragment from pENTR-FLAG-Pha21 into the 35S promoter driven pK2GW7 (57), designated pPha21-oe. For generation of A20 and/or AN1 mutant on pPha21-oe (Fig. 1), site-directed mutagenesis was conducted by QuikChange Site-Directed Mutagenesis Kit (Agilent Technologies, Santa Clara, CA, USA) and pPha21-oe was used as a template. For A20 mutant, we substituted the conserved 3rd and 4th cysteine to glycine at A20 (C35G and C38G). For AN1 mutant, we substituted the conserved 3rd cysteine and 1st histidine to glycine at AN1 (C113G and H123G). The A20 mutated clones, AN1 mutated clones, and the A20 and AN1 mutated clones, were designed as pPha21A20m, pPha21AN1m, and pPha21A20mAN1m, respectively. Primer pairs used for site directed mutagenesis are listed in Table 1.

### Transgenic *Phalaenopsis* orchid

For construction of overexpression vector to generate transgenic *Phalaenopsis* orchid, the FLAG-Pha21 fragment was transferred from pENTR-FLAG-Pha21 (described above) into 35S promoter driven binary vector, pH2GW7 (57), to obtain pHPha21. pHPha21 was used to generate transgenic *P. equestris* orchid using the method described by Hsing et al. (58).

### Agroinfiltration

Agroinfiltration was conducted as previously described by (51) with some modification. Briefly, *A. tumefaciens* C58C1 (pTiB6S3ΔT)^H^ competent cells were transformed with pCambia-CymMV, pB7GWIWG2, pK2GW7 and their derivatives using an electroporation system (Bio-Rad Laboratories, Hercules, CA, USA). Then, the *A. tumefaciens* strains were incubated at 28°C until the optical density, OD_600_, reached 0.8–1.0. After centrifugation, cells were resuspended in 20 ml AB-MES medium (17.2 mM K_2_HPO_4_, 8.3 mM NaH_2_PO_4_, 18.7 mM NH_4_Cl, 2 mM KCl, 1.25 mM MgSO_4_, 100 μM CaCl_2_, 10 μM FeSO_4_, 50 mM MES, 2% glucose (w/v), pH 5.5) with 200 μm acetosyringone (59), and cultured overnight. The overnight culture was centrifuged (3000 rpm, 10 min, in room temperature), supernatant was removed and the *A. tumefaciens* culture was resuspended in 2 ml of infiltration medium containing 50% MS medium (1/2 MS salt supplemented with 0.5% sucrose (w/v), pH 5.5), 50% AB-MES and 200 μm acetosyringone (59). The infiltration medium containing the transformed *A. tumefaciens* was applied for infiltration.

### *Cymbidium mosaic virus* (CymMV) and *Cucumber mosaic virus* (CMV) inoculation and accumulation assay

To assay the effect of Pha21, PhaDCL4, or PhaAGO1 in CymMV accumulation, we first inoculated CymMV in *P. aphrodite* through the infiltration of *A. tumefaciens* carrying pCambia-CymMV (described above) in the leaf tip of *P. aphrodite*. The CymMV-infected *P. aphrodite* was maintained at least 14 days before further analysis. To assay the effect of transient silencing (Pha21, PhaDCL4, or PhaAGO1) or transient overexpression (Pha21, or the derived mutants), *A. tumefaciens* carrying the control vector pB7GWIWG2 (for silencing), pK2GW7 (for overexpression), silencing vectors or overexpression vectors (described above) were infiltrated into the leaves. After agroinfiltration, a pair of disks (6 mm diameter) were immediately (defined as 0 dpi) collected from both the control and assay vector infiltrated regions. After 5 dpi, another pair of disks were collected from the same infiltrated region. Total RNA extracted from the samples was used as a template to analyze the accumulation of CymMV by use of qRT-PCR. The fold change of CymMV accumulation at 0 dpi to 5 dpi was calculated for relative quantification.

For inoculation with CMV, CMV-infected *Arabidopsis* leaves were ground with 0.01 M potassium phosphate buffer by pestle and mortar for use as the inoculation source. Four-week-old *Arabidopsis* leaves were inoculated mechanically (pre-dusted with 300-mesh Carborundum) with the CMV inoculation source. After 14 dpi, the disease symptoms were observed, and three disks from three different distal leaves were collected for CMV accumulation analysis by use of qRT-PCR. Primers used for CMV detection are listed in Table 1.

### Preparation and transfection of protoplasts

For construction of vectors used for subcellular localization analysis, the primer pairs, Pha21-ORF-F/Pha21-ORF-R and Pha21-ORF-F/Pha21-ORF-non-stop-R (Table 1) were used to amplify 2 sets of Pha21 ORF (with or without a stop codon). The amplified fragment was cloned into Gateway entry vector pENTR/D-TOPO (Invitrogen) to generate pENTR-Pha21 and pENTR-Pha21-non-stop following the manufacturer’s protocol. Subsequently, LR Gateway cloning reaction (Invitrogen) was performed to transfer ORF fragment of *Pha21* from pENTR-Pha21 into p2FGW7 driven by 35S promoter (57) to obtain N-terminal GFP fused clones (pG-Pha21). To obtain C-terminal GFP-fused clones (pPha21-G), we transferred and pENTR-Pha21-non-stop into p2GWF7.

Protoplast isolation and transfection were as described by Lu et al. (51). Transformed protoplasts were detected for florescence signals by confocal microscopy (Zeiss LSM 780, plus ELYRA S.1) with excitation at 488 nm and emission at 500 to 587 nm for GFP, and excitation at 543 nm and emission at 600 to 630 nm for mCherry.

### Transcriptional activation ability assay

The Pha13 and Pha21 ORF was amplified by PCR with gene specific primer pairs, NdeI-Pha13-F/EcoR1-Pha13-R and NdeI-Pha21-F/BamH1-Pha21-R, respectively (Table 1). The amplified ORF fragment was then cloned into the pGBKT7 vector (Takara Bio) to generate the pGBKT7-Pha13 and pGBKT7-Pha21 constructs, which fused to Gal4 BD sequence. These constructs were individually transformed into AH109 yeast strain (Takara Bio) and the self-activation ability was analyzed. The transformation and selection procedure was performed following the Yeast Protocols HandBook (Takara Bio). Transformants were selected on SD/-Trp (tryptophan) medium. The expression of reporter genes in yeast were tested on different selection media; SD/-Trp/Aureobasidin was used to test for activation of the AUR1-C (inositol phosphoryl ceramide synthase) reporter against Aureobasidin A, SD/-Trp/X-α-Gal medium was used to test for activation of α-galactosidase, and SD/-Trp/-Ade/-His medium was used to test for induction of the ADE2 and HIS3 reporters. The yeast containing Gal4 DNA-BD fused with murine p53 (pGBKT7-53) and the Gal4 AD fused with SV40 large T-antigen (pGADT7-T) from the Matchmaker Gold Yeast Two-Hybrid System kit (Takara Bio) were used as positive control. The yeast containing Gal4 DNA-BD fused with lamin (pGBKT7-Lam) and the pGADT7-T from the Matchmaker Gold Yeast Two-Hybrid System kit (Takara Bio) were used as a negative control.

### Construction, expression, and purification of recombinant proteins

Full-length Pha21, or Pha21 A20 and/or AN1 domain mutant fragments were amplified by PCR with primer pairs NdeI-Pha21-F/NotI-Pha21-R (Table 1) using pPha21-oe, pPha21A20m, pPha21AN1m, and pPha21A20mAN1m as templates. The amplified fragments were cloned into the pET24b expression vector (Merck, Darmstadt, Germany) to produce fused C-terminal histidine tag (His-tag) expression plasmids, pETPha21, pETPha21A20m, pETPha21AN1m and pETPha21A20mAN1m. The constructed plasmids were transformed into *Escherichia coli* strain BL21 (DE3) for protein expression. Bacteria were cultured at 37°C to an OD600 of 0.5, then transferred to 25°C, for 1.5 h. Then, isopropylthio-β-galactoside (IPTG; Sigma) was added to a final concentration of 1 mM for protein induction. His-tagged recombinant protein was purified by TALON Superflow (GE Healthcare Life Sciences, Pittsburgh, PA, USA) according to the manufacturer’s description. His-tagged recombinant protein was eluded with 250 mM imidazole (Sigma).

### E3 ubiquitin ligase activity assay

*In vitro* ubiquitination assays were performed as described by (45) with modification. An amount of 3 μg purified His-tagged recombinant proteins (*Pha21* or derived mutants) was used for each ubiquitination reaction. Reactions were incubated at 30°C for 3 hours and analyzed by SDS-PAGE followed by immunoblot analysis. Blots were probed using anti-FLAG antibodies (Sigma) followed by HRP conjugated goat anti-mouse antibodies (GE Healthcare Life Sciences).

### Yeast two-hybrid assay

For the ubiquitin binding ability assay, the pGBKT7-Pha21 (described above) was used as a bait vector. To map the region of Pha21 for ubiquitin binding, a deletion mutant of Pha21 was generated by PCR amplification with the primer pairs described in Table 1. The PCR-amplified fragment was individually cloned to pGBKT7 and used as a bait vector (Fig. 6B). Full length of orchid ubiquitin was amplified by RT-PCR with the primer pair, NdeI-PhaUBQ-F/BamHI-PhaUBQ-R, and cloned into pGADT7 vector (Takara Bio) to generate pGADT7-UBQ as a prey vector. pGBKT7-Pha21 and pGADT7-UBQ was co-transformed into AH109 yeast strain for yeast two-hybrid assay by using the Make Your Own “Mate & Plate” Library System (Takara Bio) following the user manual. Yeast strains containing the appropriate bait and prey plasmids were cultured in liquid 2-dropout medium (leucine^-^ and tryptophan^-^) overnight. The overnight yeast culture was diluted to an OD600 of 0.06 and spotted on selection plates (containing histidine^-^, leucine^-^, tryptophan^-^ for growth assay.

For the interaction analysis between Pha21 and Pha13, full-length Pha13 fragment was amplified by PCR with primer pair NdeI-Pha13-F/EcoR1-Pha13-R (Table 1). The Pha13 fragments were cloned into pGBKT7 and pGADT7 vector to generate pGBKT7-Pha13 as the bait vector and pGADT7-Pha13 as the prey vector. The full-length of Pha21 previously used to generate the pGBKT7-Pha21 (described above) was also cloned into pGADT7 to generate pGADT7-Pha21 as prey vector. The pGBKT7-Pha21, pGBKT7-Pha13, pGADT7-Pha21, and pGADT7-Pha13 were co-transformed into AH109 yeast strain for yeast two-hybrid assay as described above. The pGBKT7-53 and pGADT7-T from the Matchmaker Gold Yeast Two-Hybrid System kit were used as a positive control.

### Accession numbers

Pha21 (PATC144963), Pha13 (PATC148746), PhaPR1 (PATC126136), PhaNPR1 (PATC135791), PhadR1 (PATC143146), PhaRdR2 (PATC124544), PhaRdR6 (PATC131836), PhaDCL2 (PATC143544), PhaDCL4 (PATC150652), PhaAGO1 (PATC157237), PhaAGO10 (PATC093469), PhaGRXC9 (PATC068819), PhaUBQ10 (PATC230548), PhaJAZ1 (PATC141437), PhaACO2 (PATC139319), AtActin (At3G18780), AtSAP5 (AT3G12630), DCL4 (AT5G20320), AGO1 (AT1G48410), OsSAP3 (LOC_Os01g56040.1), OsSAP5 (LOC_Os02g32840.1)

## ACKNOWLEDGMENTS

We thank Shu-Chen Shen from the Confocal Microscope Core Facility at Academia Sinica for assistance in confocal microscopy images and Chii-Gong Tong from the Gene Transgenic Room at Academia Sinica for orchid transformation. This work was supported by grants from Academia Sinica, Taipei, Taiwan and the Ministry of Science and Technology (105-2313-B-001-004-MY3) of Taiwan. The funders had no role in study design, data collection and interpretation, or the decision to submit the work for publication

## References

1. Pieterse CM, Leon-Reyes A, Van der Ent S, Van Wees SC. 2009. Networking by small-molecule hormones in plant immunity. Nat Chem Biol 5:308–16.

2. White R. 1979. Acetylsalicylic acid (aspirin) induces resistance to tobacco mosaic virus in tobacco. Virology 99:410–412.

3. Chivasa S, Murphy AM, Naylor M, Carr JP. 1997. Salicylic acid interferes with tobacco mosaic virus replication via a novel salicylhydroxamic acid-sensitive mechanism. The Plant Cell 9:547–557.

4. Fu ZQ, Dong X. 2013. Systemic acquired resistance: turning local infection into global defense. Annual review of plant biology 64:839–863.

5. Ausubel FM. 2005. Are innate immune signaling pathways in plants and animals conserved? Nature Immunology 6:973–979.

6. Boller T, Felix G. 2009. A renaissance of elicitors: perception of microbe-associated molecular patterns and danger signals by pattern-recognition receptors. Annual review of plant biology 60:379–406.

7. Jones JD, Dangl JL. 2006. The plant immune system. Nature 444:323–329.

8. Yang H, Gou X, He K, Xi D, Du J, Lin H, Li J. 2010. BAK1 and BKK1 in Arabidopsis thaliana confer reduced susceptibility to turnip crinkle virus. European journal of plant pathology 127:149–156.

9. Kørner CJ, Klauser D, Niehl A, Domínguez-Ferreras A, Chinchilla D, Boller T, Heinlein M, Hann DR. 2013. The immunity regulator BAK1 contributes to resistance against diverse RNA viruses. Molecular plant-microbe interactions 26:1271–1280.

10. Niehl A, Wyrsch I, Boller T, Heinlein M. 2016. Double-stranded RNAs induce a pattern-triggered immune signaling pathway in plants. New Phytologist 211:1008–1019.

11. Holmes FOJBG. 1929. Local lesions in tobacco mosaic. 87:39–55.

12. HoLMEs FOJP. 1937. Inheritance of resistance to tobacco-mosaic disease in the pepper (Capsicum sp.). 27:637–642.

13. Holmes FOJP. 1938. Inheritance of resistance to tobacco-mosaic disease in tobacco. 28.

14. Holmes FO. 1954. Inheritance of resistance to viral diseases in plants, p 1–30, Advances in virus research, vol 2. Elsevier.

15. Tsuda K, Katagiri FJCoipb. 2010. Comparing signaling mechanisms engaged in pattern-triggered and effector-triggered immunity. 13:459–465.

16. Ross AFJV. 1961. Localized acquired resistance to plant virus infection in hypersensitive hosts. 14:329–339.

17. Ross AFJV. 1961. Systemic acquired resistance induced by localized virus infections in plants. 14:340–358.

18. Ding S-W, Voinnet O. 2007. Antiviral immunity directed by small RNAs. Cell 130:413–426.

19. Zhang C, Wu Z, Li Y, Wu J. 2015. Biogenesis, function, and applications of virus-derived small RNAs in plants. Frontiers in microbiology 6:1237.

20. Sharma N, Sahu PP, Puranik S, Prasad MJMb. 2013. Recent advances in plant– virus interaction with emphasis on small interfering RNAs (siRNAs). 55:63–77.

21. Blevins T, Rajeswaran R, Shivaprasad PV, Beknazariants D, Si-Ammour A, Park H-S, Vazquez F, Robertson D, Meins Jr F, Hohn TJNar. 2006. Four plant Dicers mediate viral small RNA biogenesis and DNA virus induced silencing. 34:6233–6246.

22. Deleris A, Gallego-Bartolome J, Bao J, Kasschau KD, Carrington JC, Voinnet OJS. 2006. Hierarchical action and inhibition of plant Dicer-like proteins in antiviral defense. 313:68–71.

23. Vaucheret HJTips. 2008. Plant argonautes. 13:350–358.

24. Carbonell A, Carrington JCJCoipb. 2015. Antiviral roles of plant ARGONAUTES. 27:111–117.

25. Yu D, Fan B, MacFarlane SA, Chen Z. 2003. Analysis of the involvement of an inducible Arabidopsis RNA-dependent RNA polymerase in antiviral defense. Molecular Plant-Microbe Interactions 16:206–216.

26. Yang S-J, Carter SA, Cole AB, Cheng N-H, Nelson RS. 2004. A natural variant of a host RNA-dependent RNA polymerase is associated with increased susceptibility to viruses by Nicotiana benthamiana. Proceedings of the National Academy of Sciences of the United States of America 101:6297–6302.

27. Liu Y, Gao Q, Wu B, Ai T, Guo X. 2009. NgRDR1, an RNA-dependent RNA polymerase isolated from Nicotiana glutinosa, was involved in biotic and abiotic stresses. Plant Physiology and Biochemistry 47:359–368.

28. Li Y, Muhammad T, Wang Y, Zhang D, Crabbe MJC, Liang YJPJOB. 2018. Salicylic acid collaborates with gene silencing to tomato defense against tomato yellow leaf curl virus (TYLCV).

29. Campos L, Granell P, Tárraga S, López-Gresa P, Conejero V, Bellés JM, Rodrigo I, Lisón PJPp, biochemistry. 2014. Salicylic acid and gentisic acid induce RNA silencing-related genes and plant resistance to RNA pathogens. 77:35–43.

30. Alamillo JM, Saénz P, García JAJTPJ. 2006. Salicylic acid-mediated and RNA-silencing defense mechanisms cooperate in the restriction of systemic spread of plum pox virus in tobacco. 48:217–227.

31. Xie Z, Fan B, Chen C, Chen Z. 2001. An important role of an inducible RNA-dependent RNA polymerase in plant antiviral defense. Proceedings of the National Academy of Sciences 98:6516–6521.

32. Hunter LJR, Westwood JH, Heath G, Macaulay K, Smith AG, MacFarlane SA, Palukaitis P, Carr JP. 2013. Regulation of RNA-dependent RNA polymerase 1 and isochorismate synthase gene expression in Arabidopsis. PLoS ONE 8:e66530.

33. Zhou M, Wang W, Karapetyan S, Mwimba M, Marqués J, Buchler NE, Dong XJN. 2015. Redox rhythm reinforces the circadian clock to gate immune response. 523:472.

34. Mou Z, Fan W, Dong X. 2003. Inducers of plant systemic acquired resistance regulate NPR1 function through redox changes. Cell 113:935–944.

35. Tada Y, Spoel SH, Pajerowska-Mukhtar K, Mou Z, Song J, Wang C, Zuo J, Dong X. 2008. Plant immunity requires conformational charges of NPR1 via S-nitrosylation and thioredoxins. Science 321:952–956.

36. Herrera-Vásquez A, Salinas P, Holuigue LJFips. 2015. Salicylic acid and reactive oxygen species interplay in the transcriptional control of defense genes expression. 6:171.

37. Cao H, Bowling SA, Gordon AS, Dong X. 1994. Characterization of an Arabidopsis mutant that is nonresponsive to inducers of systemic acquired resistance. The Plant Cell 6:1583–1592.

38. Delaney T, Friedrich L, Ryals J. 1995. Arabidopsis signal transduction mutant defective in chemically and biologically induced disease resistance. Proceedings of the National Academy of Sciences 92:6602–6606.

39. Shah J, Tsui F, Klessig DF. 1997. Characterization of a s alicylic a cid-i nsensitive mutant (sai1) of Arabidopsis thaliana, identified in a selective screen utilizing the SA-inducible expression of the tms2 gene. Molecular Plant-Microbe Interactions 10:69–78.

40. Wong CE, Carson RA, Carr JP. 2002. Chemically induced virus resistance in Arabidopsis thaliana is independent of pathogenesis-related protein expression and the NPR1 gene. Molecular plant-microbe interactions 15:75–81.

41. Kachroo P, Yoshioka K, Shah J, Dooner HK, Klessig DF. 2000. Resistance to turnip crinkle virus in Arabidopsis is regulated by two host genes and is salicylic acid dependent but NPR1, ethylene, and jasmonate independent. The Plant Cell 12:677–690.

42. Vij S, Tyagi AK. 2008. A20/AN1 zinc-finger domain-containing proteins in plants and animals represent common elements in stress response. Functional & integrative genomics 8:301–307.

43. Giri J, Dansana PK, Kothari KS, Sharma G, Vij S, Tyagi AK. 2013. SAPs as novel regulators of abiotic stress response in plants. Bioessays 35:639–648.

44. Chang L, Chang H-H, Chang J-C, Lu H-C, Wang T-T, Hsu D-W, Tzean Y, Cheng A-P, Chiu Y-S, Yeh H-HJPp. 2018. Plant A20/AN1 protein serves as the important hub to mediate antiviral immunity. 14:e1007288.

45. Kang M, Fokar M, Abdelmageed H, Allen RD. 2011. Arabidopsis SAP5 functions as a positive regulator of stress responses and exhibits E3 ubiquitin ligase activity. Plant molecular biology 75:451–466.

46. Kang M, Lee S, Abdelmageed H, Reichert A, Lee HK, Fokar M, Mysore KS, Allen RD. 2017. Arabidopsis stress associated protein 9 mediates biotic and abiotic stress responsive ABA signaling via the proteasome pathway. Plant, cell & environment 40:702–716.

47. Choi H, Han S, Shin D, Lee S. 2012. Polyubiquitin recognition by AtSAP5, an A20-type zinc finger containing protein from Arabidopsis thaliana. Biochemical and biophysical research communications 419:436–440.

48. Verhelst K, Carpentier I, Kreike M, Meloni L, Verstrepen L, Kensche T, Dikic I, Beyaert R. 2012. A20 inhibits LUBAC-mediated NF-κB activation by binding linear polyubiquitin chains via its zinc finger 7. The EMBO Journal 31:3845–3855.

49. Wertz IE, O’rourke KM, Zhou H, Eby M. 2004. De-ubiquitination and ubiquitin ligase domains of A20 downregulate NF-kappaB signalling. Nature 430:694.

50. Mattera R, Tsai YC, Weissman AM, Bonifacino JS. 2006. The Rab5 guanine nucleotide exchange factor Rabex-5 binds ubiquitin (Ub) and functions as a Ub ligase through an atypical Ub interacting motif and a zinc finger domain. The Journal of Biological Chemistry 281:6874–6883.

51. Lu HC, Hsieh MH, Chen CE, Chen HH, Wang HI, Yeh HH. 2012. A high-throughput virus-induced gene-silencing vector for screening transcription factors in virus-induced plant defense response in orchid. Molecular Plant-Microbe Interactions 25:738–746.

52. Chang C-H, Wang H-I, Lu H-C, Chen C-E, Chen H-H, Yeh H-H, Tang CY. 2012. An efficient RNA interference screening strategy for gene functional analysis. BMC genomics 13:491.

53. Ma A, Malynn BA. 2012. A20: linking a complex regulator of ubiquitylation to immunity and human disease. Nature reviews Immunology 12:774.

54. Hishiya A, Iemura Si, Natsume T, Takayama S, Ikeda K, Watanabe K. 2006. A novel ubiquitin-binding protein ZNF216 functioning in muscle atrophy. The EMBO Journal 25:554–564.

55. Miyata N, Okumoto K, Mukai S, Noguchi M, Fujiki Y. 2012. AWP1/ZFAND6 Functions in Pex5 Export by Interacting with Cys-Monoubiquitinated Pex5 and Pex6 AAA ATPase. Traffic 13:168–183.

56. Giri J, Vij S, Dansana PK, Tyagi AK. 2011. Rice A20/AN1 zinc-finger containing stress-associated proteins (SAP1/11) and a receptor-like cytoplasmic kinase (OsRLCK253) interact via A20 zinc-finger and confer abiotic stress tolerance in transgenic Arabidopsis plants. New Phytologist 191:721–732.

57. Karimi M, Inzé D, Depicker A. 2002. GATEWAY™ vectors for Agrobacterium-mediated plant transformation. Trends in plant science 7:193–195.

58. Hsing H-X, Lin Y-J, Tong C-G, Li M-J, Chen Y-J, Ko S-S. 2016. Efficient and heritable transformation of Phalaenopsis orchids. Botanical studies 57:30.

59. Wu H-Y, Liu K-H, Wang Y-C, Wu J-F, Chiu W-L, Chen C-Y, Wu S-H, Sheen J, Lai E-M. 2014. AGROBEST: an efficient Agrobacterium-mediated transient expression method for versatile gene function analyses in Arabidopsis seedlings. Plant methods 10:19.

